# FAIMS-enabled N-terminomics analysis reveals novel legumain substrates in murine spleen

**DOI:** 10.1101/2023.07.18.549248

**Authors:** Alexander R. Ziegler, Antoine Dufour, Nichollas E. Scott, Laura E. Edgington-Mitchell

## Abstract

Aberrant levels of the asparaginyl endopeptidase legumain have been linked to inflammation, neurodegeneration and cancer, yet our understanding of this protease is incomplete. Systematic attempts to identify legumain substrates have previously been confined to *in vitro* studies, which fail to mirror physiological conditions and obscure biologically relevant cleavage events. Using high-field asymmetric waveform ion mobility spectrometry (FAIMS), we developed a sensitive and streamlined approach for proteome and N-terminome analyses in a single analytical method without the need for N-termini enrichment. Compared to unfractionated proteomic analysis, we demonstrate FAIMS fractionation improves neo-N- termini identification by >2.5 fold, resulting in identification of >2,882 unique neo-N-termini from limited sample amounts. Within murine spleens, this approach identifies 6,366 proteins and 2,528 unique neo-N-termini, with 235 cleavage events enriched in wild-type compared to legumain-deficient spleens. Among these, 119 neo-N-termini arose from asparaginyl endopeptidase activities, representing novel putative physiological legumain substrates. The direct cleavage of selected substrates by legumain was confirmed using *in vitro* assays, providing support for the existence of physiologically relevant extra-lysosomal legumain activity. Combined, these data shed critical light on the functions of legumain and demonstrates the utility of FAIMS as an accessible method to improve depth and quality of N- terminomics studies.

## Introduction

Proteases comprise approximately 3% of the human genome and catalyse the cleavage of peptide bonds^1^. Proteolysis is essential for maintaining protein homeostasis, altering substrate structure, function, and localisation^2^. Proteases contribute to vital cellular functions such as cell growth and repair^3^, immune signalling^4^, and wound healing^5, 6^; dysregulated protease activities underpin numerous pathological conditions including cancer, inflammation, neurodegeneration, and gastrointestinal diseases^7^. Knowledge of specific cleavage events is crucial in understanding the mechanistic contributions of proteases to normal physiology and disease. The need for sensitive approaches to catalogue these events has led to the development of peptide-centric N-terminomics methods, which have rapidly developed over the last decade^8–10^. Leveraging advancements in liquid chromatography mass spectrometry (LC-MS), peptide-based N-terminomics methods have become the gold standard for identification of protease substrates at scale^1^^1^. These techniques provide site specific resolution of cleavage events and have led to substrate discovery for numerous proteases including matrix metalloproteases (MMP) MMP-2 and MMP-9^1^^2^, caspases^13, 14^, ADAMTS7^15^, HTRA1^16^ and cathepsins^17, 18^. While N-terminomics techniques have improved the ability to identify biologically important cleavage events, these approaches are not without their limitations.

Current generation N-terminomics methods typically involve the enrichment of N-terminal peptides using either positive or negative enrichment methods^19^. These methods uniformly involve tagging the N-terminal α-amines of proteins prior to *in vitro* proteolytic digestion, therefore allowing native N-termini, which contain a defined chemical tag or native acetylation event, to be differentiated from the unmodified, newly generated N-termini (neo-N-termini)^9^. The tagging of α-amines with enrichable chemical handles permits effective positive selection of neo-N-termini, including chemical labelling with biotin^20–23^ or phosphotags^24^, or enzyme-mediated conjugates (e.g., subtiligase)^25, 26^. In contrast, negative selection approaches leverage the newly exposed α-amines of N-termini to allow the depletion of internal peptides using either N-hydroxysuccinimide polymers or resins, such as those used in TAILS (Terminal Amine Isotopic Labelling of Substrates)^27, 28^ or Nrich^29, 30^. Alternatively, internal peptides may be hydrophobically tagged as in HYTANE^31^ and HUNTER^32^, or chromatographically separated as in the COFRADIC method^33, 34^, to allow peptide depletion. While both positive and negative selection have proven effective to identify new protease substrates, these enrichment-based approaches have typically required large sample inputs, expensive and specialised materials, and user expertise, all while sacrificing the acquisition of total proteomic information. While multiple N-terminomics studies have sought to address this issue by examining non-enriched samples in parallel^18, 35–37^, the depth of these proteomic analyses is typically limited with many of the observed N-termini not quantified at the protein level. Thus, while powerful, current N-terminomics approaches still provide limited proteomic depth and utilise technologies not accessible to the broader protease and proteomics community.

A widely utilised approach to improve proteome depth is the use of orthogonal chromatographic fractionation prior to LC-MS^38, 39^. While a range of chromatographic approaches exist to fractionate proteomics samples, an alternative and increasingly accessible technology is the use of high-field asymmetric waveform ion mobility spectrometry (FAIMS)^40, 41^. This technology allows gas phase-based fractionation of peptides following chromatographic separation by filtering ion populations prior to their introduction into the mass spectrometer^42^. FAIMS allows for the fractionation of samples without the need for off- line sample handling, which is cumbersome and leads to significant sample loss^43^. Moreover, FAIMS is uniquely suited for limited sample amounts, allowing enhanced detection sensitivity^44^ and dramatically improved proteomic depth^45^. Whilst widely used to improve proteome coverage, FAIMS-based approaches have also been shown to dramatically improve the identification of peptide subsets including cross-linked peptides^46^, cysteine-containing peptides^47^ and glycopeptides^48^. Inspired by these previous studies, we set out to assess whether FAIMS-based analysis would allow for deep proteomic coverage and simultaneous assessment of the N-terminome on limited samples, yielding a cheaper and more streamlined method to identify protease substrates, using the protease legumain as a model system.

Legumain is a cysteine protease with unique preference to cleave substrates after asparagine residues^49^. Following synthesis as an inactive zymogen, it is trafficked to the endo-lysosomal pathway via mannose-6-phosphate-dependent mechanisms. Upon reaching acidic environments, legumain cleaves itself to produce a mature, proteolytically active enzyme^50, 51^. Legumain protease activity favours acidic conditions, and its active conformer is thought to be rapidly destroyed upon entering neutral environments^51^. Increasing evidence suggests extra- lysosomal localisation of legumain, and that it can cleave substrates in these environments; including the nucleus^52^, cytoplasm^53^, and extracellular space^54^. While binding to integrins via its RGD motif is postulated to stabilize active legumain at the cell surface, how it remains active in the nucleus and cytoplasm is not well understood.

Legumain contributes to renal homeostasis and lysosomal protein turnover, as evidenced by renal insufficiency and lysosomal storage disorders in legumain-deficient mice^55–57^. Legumain activity is upregulated in a range of diseases, including Alzheimer’s and Parkinson’s diseases^58–59, 60^, pancreatitis^61, 62^, and cancer^63, 64^. Inhibiting legumain reduced synapse loss and cognitive impairment in tauopathy mice^65, 66^ and reduced α-synuclein cleavage in SNCA-transgenic mice to improve dopamine levels and motor functions^67^. In a MMTV-PyMT murine breast cancer model, blocking legumain activity decreased lung metastasis^63^. These studies suggest that targeting legumain activity has strong therapeutic potential. The proteolytic events leading to these observed phenotypes, however, are yet to be fully elucidated. To date, relatively few legumain substrates have been identified^49^, among which include the invariant chain^68, 69^, pro- MMP-2^70^, endosomal toll-like receptors^71^, and the nuclear protein FOXP3^72^. Recent studies have aimed to systematically identify legumain substrates by spiking recombinant legumain into acidified lysates^73, 74^. In these *in vitro* conditions, legumain cleaves hundreds of proteins after asparagine residues, and at lower pH, also after aspartate residues. Identification of physiological substrates, where cellular compartmentalisation and pH environments are intact, however, is lacking. To better understand the proteolytic contribution of legumain to cellular function and disease, an unbiased and systematic approach to identify its native substrates is required.

In the current study, we benchmarked our FAIMS-enabled N-terminomics method in mouse macrophages treated with the legumain inhibitor SD-134^75^, revealing significant improvements in coverage of N-termini compared to unfractionated samples. We then analysed naïve spleens from wild-type and legumain-deficient (*Lgmn^-/-^*) mice to reveal global alterations in proteolysis, including 119 putative legumain substrates. Our data provide the first comprehensive list of physiological legumain substrates which provides insight into novel functions of legumain in neutral cellular environments. FAIMS-enabled N-terminomics is thus a streamlined and accessible method to identify novel proteolytic substrates.

## Results

### FAIMS fractionation enables deep proteome coverage and identification of native cleavage events

Conventional N-terminomics techniques often rely on selection methods to enrich N-terminal peptides^19^. Whilst effective, enrichment is performed at the cost of bulk proteome data, limiting the ability to assess if observed alterations are true changes in the N-termini or the global proteome. To overcome this, we assessed the potential to undertake simultaneous proteome and N-terminome analyses by coupling FAIMS fractionation to established dimethylation-based N-termini labelling^27, 32^. Subsequent fragment-ion indexing-based proteomic searches using MSFragger^76^ allowed the searching of datasets with large combinations of variable modifications (**Fig. 1a**). We reasoned this would enable identification and quantitation of both protein abundance and cleavage event (neo-N-termini) information from limited amounts of complex samples such as tissue.

**Fig. 1.**
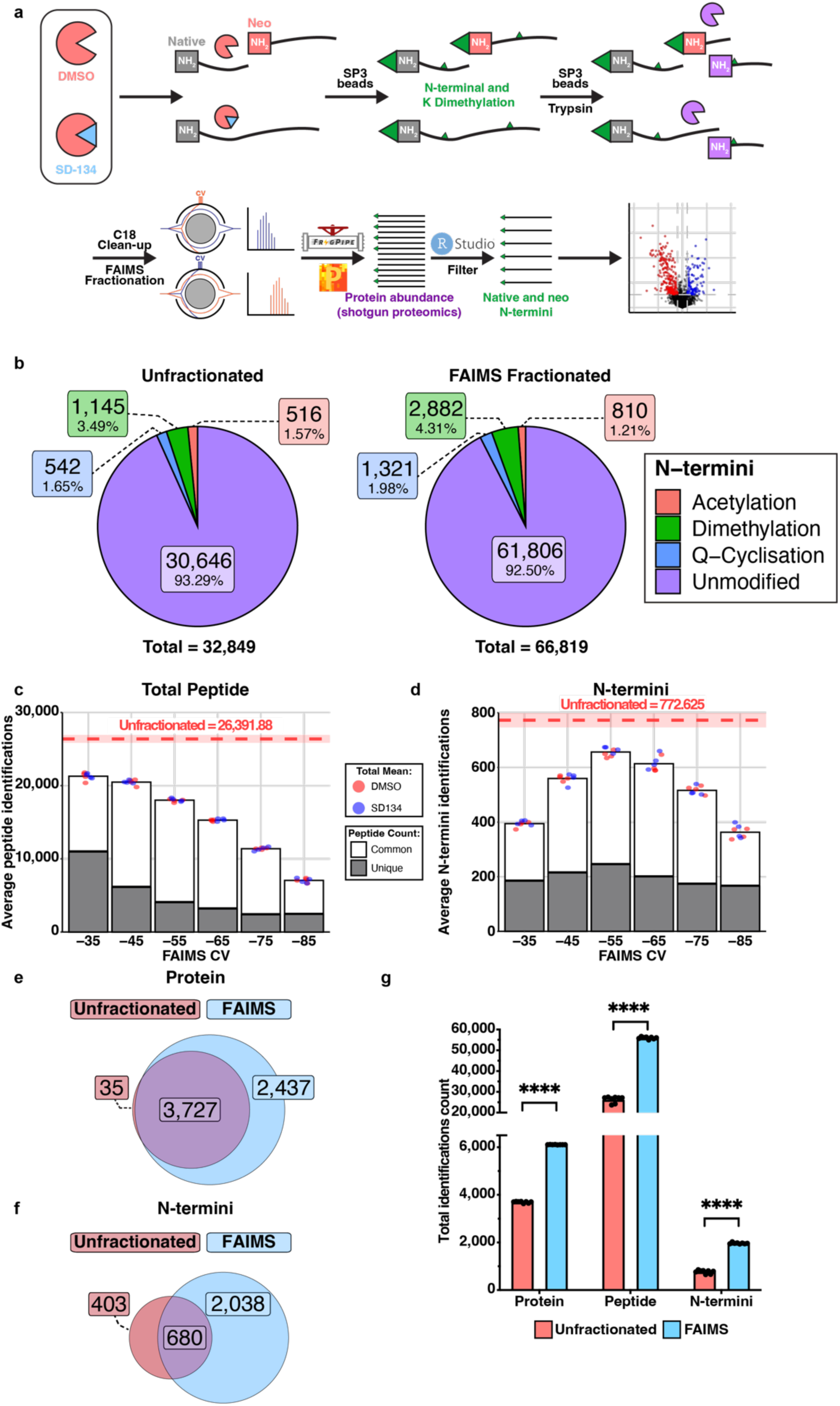
FAIMS-enabled N-terminomics increases overall peptide detection compared to unfractionated methods. a. Experimental workflow. RAW264.7 cells treated with DMSO (n = 4) or 10 µM SD-134 (n = 4) and naïve spleen tissue from wild-type (n = 4) and legumain- deficient (*Lgmn-/-*, n = 4) mice were analysed by FAIMS-enabled N-terminomics. Native and neo-N-termini were labelled with formaldehyde and peptides digested by trypsin. Online gas- phase fractionation was achieved using FAIMS (high-field asymmetric wavefield ion mobility spectrometry) over a range of compensation voltages (CV, -35, -45, -55, -65, -75, -85) prior to mass spectrometry analysis. Data were analysed by MSFragger (Fragpipe v.18.0) and Perseus (v.1.6.0.7). Native cleavage sites were bioinformatically enriched by filtering for N-terminal dimethylation using RStudio. Numbers shown refer to peptide-spectrum matches present in at least three of four biological replicates in minimum one group (**b-f**). **b.** Peptide-spectrum matches, and their N-terminal modifications identified in unfractionated (left panel) and FAIMS-fractionated (right panel) RAW264.7 cell lysates. **c-d.** Average total peptides (**c**) and dimethylated N-terminal peptides (neo-N-termini) (**d**) identified in each biological replicate per FAIMS fraction (CV=compensation voltage). Red dashed line refers to average peptides or N-termini identified without FAIMS fractionation with standard deviation indicated by the box . Peptide-spectrum matches identified in only one specified CV fraction are indicated in grey (unique to that fraction) and those identified in multiple CV fractions are shown in white (common between fractions). **e-f.** Unique protein (**e**) and neo-N-termini (**f**) identifications and their overlap (purple) between unfractionated (red) and FAIMS-fractionated (blue) RAW264.7 cell lysates. **g.** Average number of proteins, peptides, and neo-N-termini identified in each biological replicate from unfractionated (red) and FAIMS-fractionated (blue) RAW264.7 cell lysates. A student’s t-test was performed for pairwise comparisons. (****p<0.0001).

We applied FAIMS to analyse the global proteome and N-terminome of RAW264.7 murine macrophages in response to the legumain-specific inhibitor SD-134^75^. Using the fluorescently quenched activity-based probe for legumain, LE28, we confirmed the ability of SD-134 to inhibit legumain^77^ (**Supplementary Fig. 1**). To investigate whether FAIMS could improve N-terminome analysis, protein lysates were dimethylated, digested with trypsin and subjected to online gas-phase fractionation using six FAIMS compensation voltages (CVs: -35, -45, -55, - 65, -75 and -85) (**Supplementary Table 1, 2, 6 and 7**), with each analytical run utilising 2 μg digested material. This was benchmarked against identical analysis in the absence of FAIMS fractionation. A total of 32,849 unique peptides corresponding to 3,762 proteins were identified in unfractionated samples, while FAIMS permitted detection of 66,819 peptides corresponding to 6,164 proteins (2.03-fold increase) (**Fig. 1b, Supplementary Table 3, 4, 8 and 9**). The dimethylation labelling efficacy was observed to be >95% (**Supplementary Fig. 2**). Whilst the percentage of dimethylated N-termini observed with and without FAIMS fractionation were similar (3.49% unfractionated and 4.31% fractionated of total peptide, **Fig. 1b**), FAIMS provided access to a greater number of neo-N-termini (2,882) than unfractionated samples (1,145) (**Supplementary Table 5, 10 and 11**). Furthermore, as each FAIMS fraction yielded a substantial number of peptide and neo-N-termini identifications exclusive to that CV value (**Fig. 1c-d**), coverage of the N-terminome was improved.

Whilst the majority of protein and neo-N-termini identified within the unfractionated samples were also identified after FAIMS fractionation, the improved proteome depth of this approach lead to detection of an additional 2,038 unique dimethylated peptides (**Fig. 1e-f**). Of the quantified neo-N-termini, approximately 95% had complete protein quantifications in at least one of the biological groups, which was marginally elevated compared to unfractionated samples (90%) (**Supplementary Fig. 3**). Overall, FAIMS fractionation identified significantly more proteins, peptides, and neo-N-termini per biological replicate compared to unfractionated samples (**Fig. 1g**). We observed tighter distributions and significant reductions in the standard deviation of protein and neo-N-termini quantifications following FAIMS fractionation, demonstrating that the quality of the data was also improved (**Supplementary Fig. 4a-d**). When comparing our results to a previously published TAILS dataset obtained from RAW264.7 cells treated with the legumain inhibitor LI-1^78^ (**Supplementary Fig. 5**), we detected a similar number of neo-N-termini in the two methods, despite using >10x the amount of starting material for the TAILS analysis. Hence, FAIMS-enabled N-terminomics circumvents the requirement of N-termini enrichment, allowing assessment of both the proteome and N-terminome using limited sample amounts.

Compared to unfractionated analysis, FAIMS fractionation also produced an expanded set of proteins and neo-N-termini exhibiting significant differences between DMSO- and SD-134- treated samples (**Fig. 2a, b, d, e; Supplementary Table 3, 5, 8 and 10**). We hypothesise that greater proteome coverage (**Fig. 1b**) and the reduced variance in the data distribution observed with FAIMS (**Supplementary Fig. 4**) may account for these differences. To identify legumain-specific cleavage events, we filtered the dimethylated N-termini for those that arose due to cleavage after asparagine residues. While no asparaginyl cleavages were significantly enriched in unfractionated DMSO-treated samples (**Fig. 2c; Supplementary Table 5**); five were detected by the FAIMS method (**Fig. 2f; Supplementary Table 10-11**). One of these cleavage sites was within another lysosomal protease, cathepsin S; the neo-N-termini identified corresponded to cleavage at N^120^↓R^121^, immediately upstream of the canonical pro-cathepsin S cleavage site (**Fig. 2g**). We validated the direct cleavage of cathepsin S by legumain *in vitro* using recombinant human proteins, demonstrating processing of cathepsin S from its 37 kDa proform to a 25 kDa mature form (**Fig. 2h**). Using unfractionated N-terminomics analysis, we demonstrated that legumain cleaves human cathepsin S *in vitro* at the conserved site N^112^ ↓ R^113^ (**Fig. 2i**, **Supplementary Fig. 6; Supplementary Table 12**). Together, our data demonstrate that FAIMS-enabled N-terminomics is an effective workflow for the sensitive detection of neo-N-termini from complex samples, allowing protease substrate identification without enrichment.

**Fig. 2.**
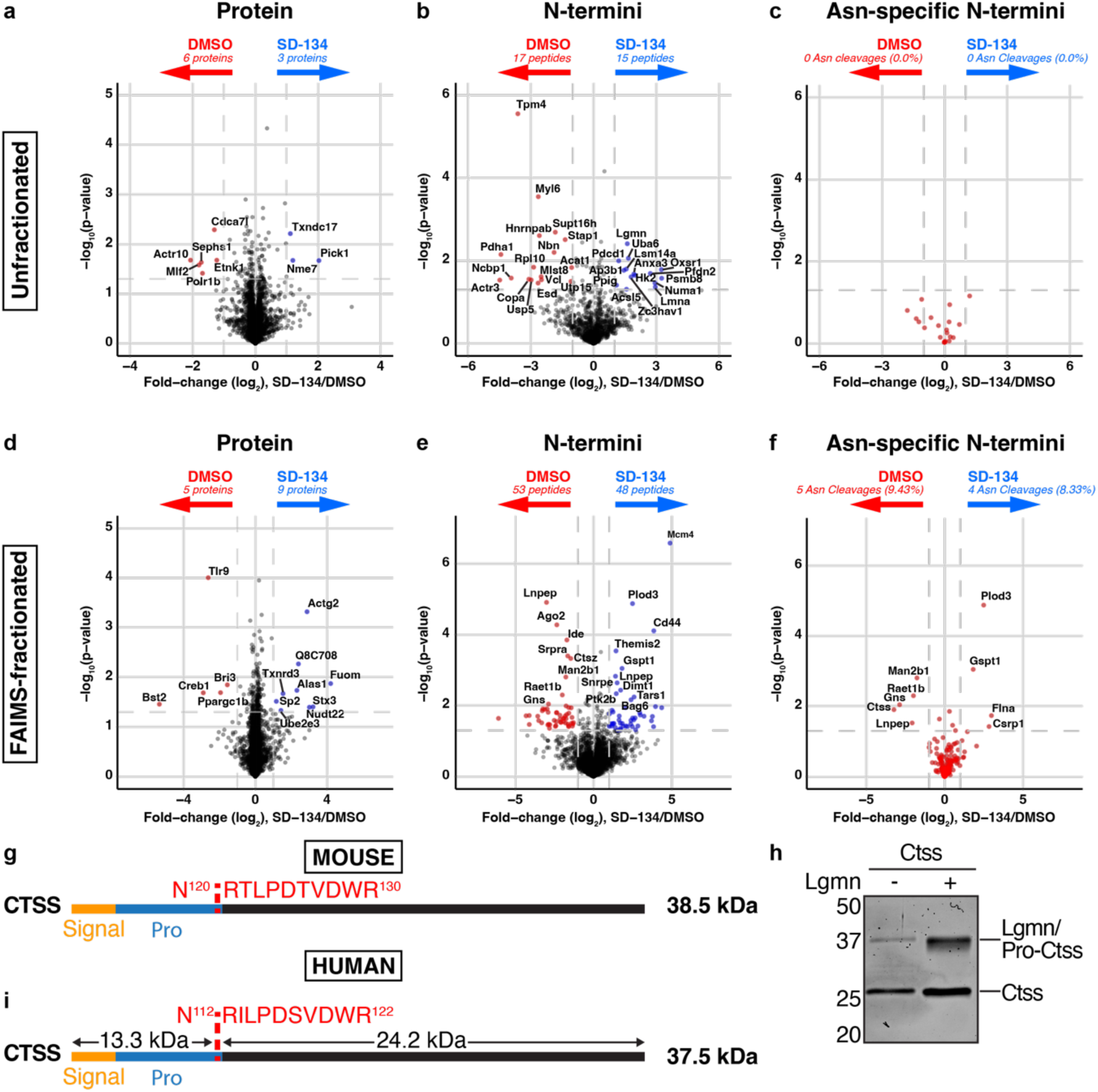
Quantitative proteomics and N-terminomics analyses of unfractionated and FAIMS- fractionated DMSO and SD-134-treated RAW 264.7 cell lysates. Peptide-spectrum matches were analysed using Perseus and filtered to have valid values within at least three of four biological replicates in at least one group. **a-f.** For unfractionated (**a-c**) and FAIMS-fractionated (**d-f**) data, protein (**a, d)** and neo-N-termini (**b, e**) identifications were analysed by a two-way t-test and visualised by volcano plot where significance is defined as abs(log2(SD-134/DMSO)) > 1 and -log10(p) > 1.3. Neo-N-termini arising from cleavage after asparagine residues were identified and highlighted in red (**c, f**). **g.** Schematic of murine cathepsin S (Ctss) cleavage at N120 ↓ R121 as identified by FAIMS-enabled N-terminomics. Cleavage site is shown in red, signal peptide in orange, and propeptide in blue. **h.** SYPRO Ruby-stained gel of recombinant human cathepsin S (CTSS) incubated with (+) or without (-) activated recombinant human legumain (LGMN) at pH 5.5 for 5 hours (1:1 mass ratio). Image is representative of 3 independent experiments with similar results. **i.** Schematic representation of human cathepsin S cleavage by legumain at N112 ↓ R113 as identified by N-terminomics analysis of recombinant protein cleavage assay. Predicted size of resulting cleavage products are shown. Identified cleavage site is shown in red, signal peptide in orange, and propeptide in blue.

### FAIMS-enabled analysis of legumain-deficient mouse spleens reveals altered proteolysis and neutrophil function

We next aimed to apply FAIMS-enabled N-terminomics to examine the influence of legumain on the global proteome and N-terminome in a more physiologically relevant setting. As relatively little information is available on the role of legumain in spleen, we analysed naïve spleens from wild-type (WT) and legumain-deficient (*Lgmn^-/-^*) mice (**Supplementary Table 13- 14**). We confirmed legumain was present and active in WT splenic lysates and absent in *Lgmn^-/-^* lysates using the LE28 activity-based probe^77^ and immunoblot (**Supplementary Fig. 7**); this was further verified in our LC-MS/MS analysis (**Fig. 3a**). We identified 64,649 peptides from 6,366 proteins in our FAIMS-fractionated spleen lysates across all biological replicates (**Supplementary Table 15-16**), where samples demonstrated clear clustering based on biological groups (**Supplementary Fig. 8**). Among the detected peptides, 2,528 were dimethylated, with a labelling efficacy >95% (**Supplementary Table 17; Supplementary Fig. 9)**. Additionally, each FAIMS fraction revealed a unique set of N-terminal peptides enabling deep N-terminome coverage (**Supplementary Fig. 10**).

**Fig. 3.**
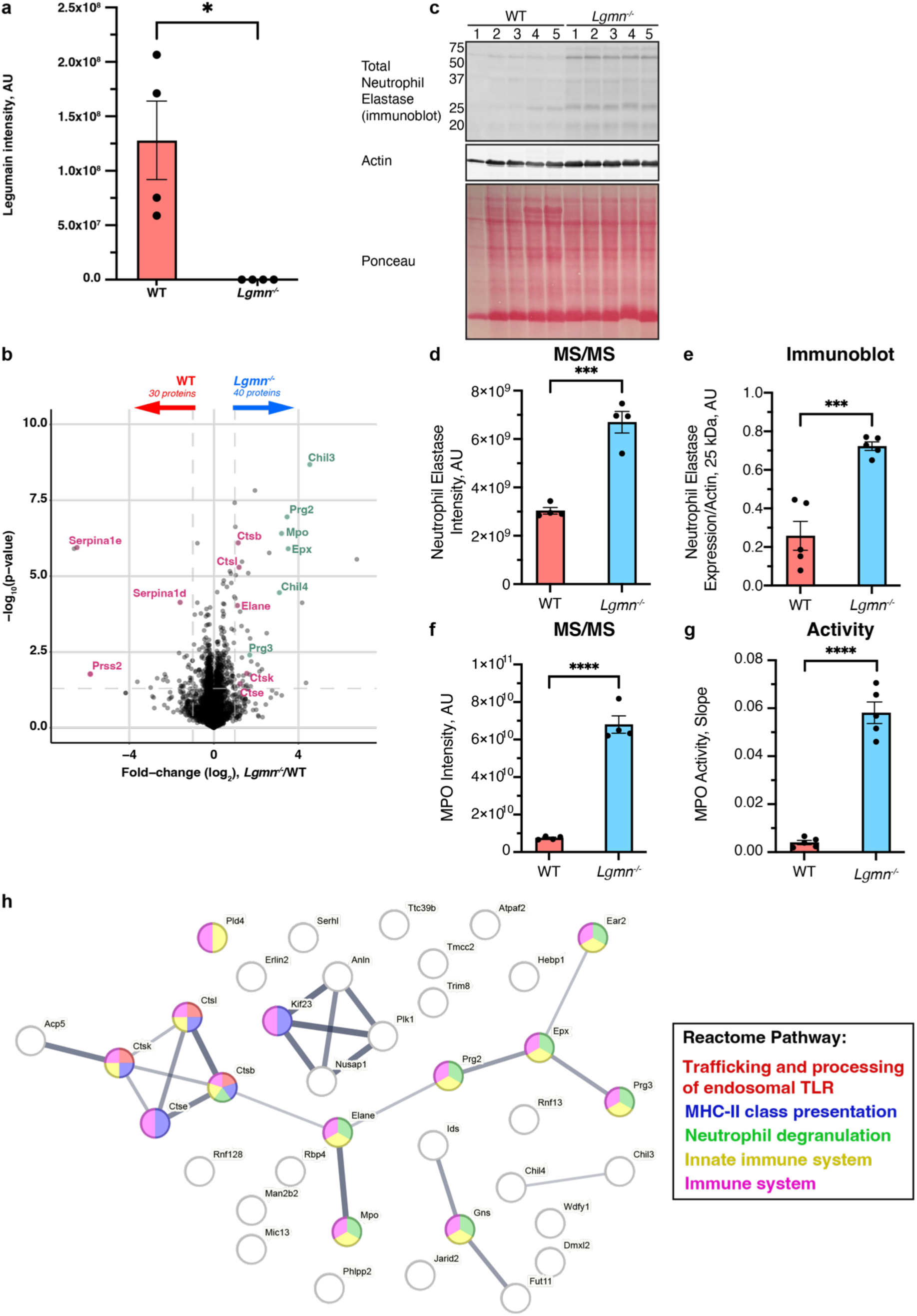
Quantitative proteomics analysis of wild-type (WT) and legumain knockout (*Lgmn-/-*) naïve mouse spleens by FAIMS-enabled N-terminomics. a. Legumain intensity values from quantitative proteomics analysis of WT and *Lgmn-/-* naïve mouse spleens. Biological replicates are shown. A student’s t-test was performed for pairwise comparisons (*p < 0.05). Proteins identified in ≥3 of 4 biological replicates in at least one group (n = 4/group) were analysed by a two-way t-test and visualised by volcano plot (**b**). Significantly elevated proteins are defined by abs(log2(*Lgmn^-/-^*/WT)) > 1 and -log10(p) > 1.3. Altered proteins related to proteolysis are shown in magenta and those corresponding to neutrophils in green. **c.** Total neutrophil elastase expression in naïve mouse spleen lysates measured by immunoblot. Actin and Ponceau S stain were used as loading controls. **d.** Neutrophil elastase intensity measurements from proteomic analysis of WT and *Lgmn-/-* naïve mouse spleens (n = 4/group). **e.** Immunoblot bands were quantified by densitometry and normalised to actin (n = 5/group). **f.** Myeloperoxidase intensity measurements from proteomic analysis of WT and *Lgmn-/-* naïve mouse spleens (n = 4/group). **g.** Mouse spleen lysates were analysed by a myeloperoxidase activity assay and linear slope was calculated from 0-6 minutes (n = 5/group). A student’s t- test was performed for pairwise comparisons (ns = not significant, *p < 0.05, **p < 0.01, ***p < 0.001). **h.** STRING-db (v.11.5) analysis of the 40 *Lgmn-/-* enriched proteins (confidence = 0.400, false discovery rate = 5%). Line thickness corresponds to the confidence of interaction. Reactome pathway: red=trafficking and processing of endosomal TLR, blue=MHC-II class presentation, green=neutrophil degranulation, yellow=innate immune system, magenta=immune system.

We observed 30 proteins that were reduced in abundance upon loss of legumain, including trypsin-2 (*Prss2)* and serine protease inhibitors (*Serpina1e* and *Serpina1d*) (**Fig. 3b**). Conversely, 40 proteins exhibited increased abundance in the absence of legumain including cathepsin B, L, E, and K, and several neutrophil-associated proteins (neutrophil elastase, myeloperoxidase, eosinophil peroxidase, proteoglycans, and chitinase-like proteins) (**Fig. 3b, Supplementary Fig. 11**). We verified an increase in neutrophil elastase in *Lgmn^-/-^* spleens by immunoblot (**Fig. 3c- e**). Likewise, myeloperoxidase activity was dramatically amplified in *Lgmn^-/-^* spleens (**Fig. 3f-g**). STRING (v.11.5) analysis of the 40 *Lgmn^-/-^*-enriched proteins further revealed alterations in immune-related pathways including toll-like receptor (TLR) processing, MHC-II class presentation, and neutrophil degranulation (**Fig. 3h**). Consistently, proteins contributing to these pathways were overexpressed in *Lgmn^-/-^* spleen lysates compared to WT counterparts (**Supplementary Fig. 12**). Overall, our proteomics data suggest the involvement of legumain in proteolytic regulation and neutrophil function.

### Identification of novel legumain substrates within naïve mouse spleens

To identify legumain-mediated alterations of the N-terminome, we compared the dimethylated N-termini observed between WT and *Lgmn^-/-^* naïve mouse spleens. Of 2,528 neo-N-termini (**Fig. 4a, Supplementary Table 17**), 1,443 of these were not within the first 65 amino acids of the proteins, consistent with endopeptidase activity (**Supplementary Fig. 13a- c**). Statistical analysis (log2(fold change) > ±1 and -log10(p) > 1.3) revealed 235 cleavage events enriched in WT spleens and 116 in *Lgmn^-/-^* (**Fig. 5b, Supplementary Table 17**). Amino acids flanking the observed cleavage sites were assessed to examine cleavage motifs enriched in WT and *Lgmn^-/-^* tissue (**Fig. 4e, 4f**). In line with the well-established asparaginyl endopeptidase activity of legumain, WT samples exhibited a strong enrichment in neo-N-termini arising from cleavage after asparagine (Benj. Hoch. FDR = 1.14e^-41^, **Supplementary Table 18-19**). In fact, 50.6% (119/235) of the WT-enriched neo-N-termini were cleaved after Asn, while this was observed in only 2.6% (3/116) of the neo-N-termini from *Lgmn^-/-^* spleens (**Fig. 4c, 4d; Supplementary Table 19-20**).

**Fig. 4.**
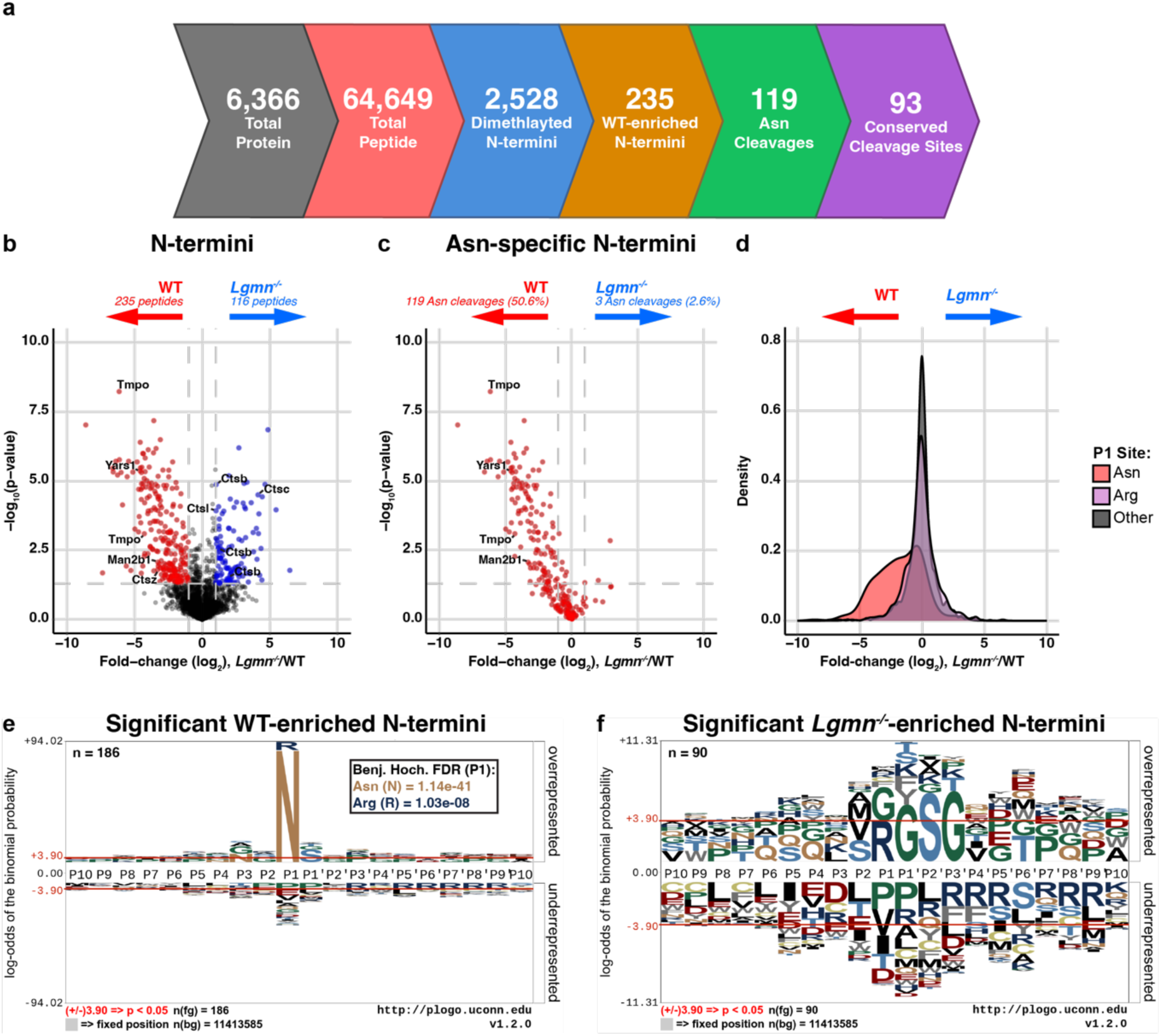
FAIMS-enabled N-terminomics analysis of wild-type (WT) and legumain-deficient (*Lgmn-/-*) naïve mouse spleens. Identified peptides were filtered for those present in at least 3 of 4 biological replicates in at least one group (n = 4/group). **a.** Proteins and peptides identified in FAIMS-fractionated naïve mouse spleen lysates are summarised. Total peptide- spectrum matches were bioinformatically filtered for N-terminal dimethylation indicating endogenous N-termini. Neo-N-termini were further filtered for those arising due to cleavage after asparagine residues, and for those that are conserved in both mouse and human proteins. A two-sample t-test was performed and N-termini were visualised by volcano plot (**b-c**). **b.** WT-enriched neo-N-termini are shown in red (log2(*Lgmn^-/-^*/WT) < -1 and -log10(p) > 1.3) and *Lgmn^-/-^* in blue (log2(*Lgmn^-/-^*/WT) > 1 and -log10(p) > 1.3). **c.** Asparaginyl cleavage events were identified and highlighted in red. **d.** Density plot of data shown in Fig. 5b showing log2(*Lgmn^-/-^*/WT) distribution of neo-N-termini. Peptides arising due to cleavage after asparagine residues are shown in red, arginine residues in purple and all other residues in grey. **e-f**. Sequence motifs of neo-N-termini significantly enriched in WT (n = 186) (**e**) and *Lgmn-/-* (n = 90) (**f**) naïve spleen lysates were created using plogo (O’Shea et al. 2013). Overrepresented amino acids appear above and underrepresented below the x-axis (p < 0.05).

**Fig. 5.**
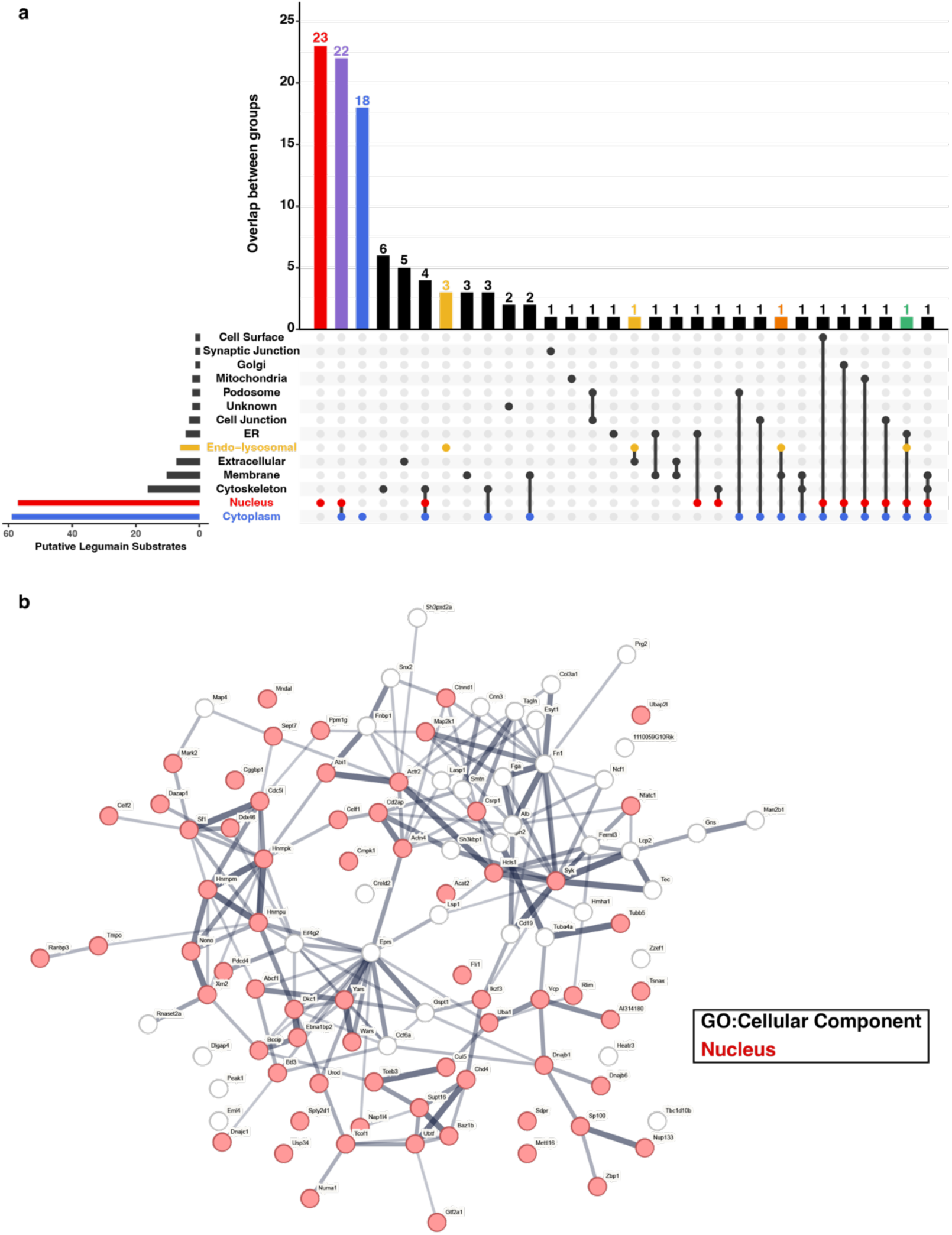
Identification of putative legumain substrates in naïve mouse spleen lysates and characterisation of their subcellular localisation. Legumain substrates were classified as neo- N-termini significantly enriched in WT naïve spleen lysates (log2(*Lgmn^-/-^*/WT) < -1 and -log10(p) > 1.3) containing an asparagine residue in the P1 site. **a.** Upset plot of subcellular localisations of the 119 putative legumain substrates identified. Information is taken from UniProt (https://www.uniprot.org/). Compartments of interest are highlighted such that red indicates nucleus, blue indicates cytoplasm, and yellow indicates endo-lysosomal system. Proteins localised to both nuclear and cytoplasmic regions are highlighted in purple, both cytoplasmic and endo-lysosomal in orange, and all three nuclear, cytoplasmic, endo-lysosomal in green. **b.** Putative legumain substates were further analysed by STRING-db for gene ontology terms, cellular component (GO:CC). Red indicates nuclear localisation of the protein.

Among the WT-enriched P1 asparaginyl cleavages (**Supplementary Fig. 13d; Supplementary Table 21**), we observed slight preferences for serine in the P1’ position, proline in the P3’, and glycine in the P2 and P3 positions. We also visualised the consensus motif for the non- asparaginyl cleavages (**Supplementary Fig. 13e; Supplementary Table 22**). Despite the well-established ability of legumain to cleave after aspartate residues in acidic pH, P1 Asp was only observed in 4 of the 116 non-asparaginyl cleavages. Notably, asparagine in the P3 position was enriched among these neo-N-termini (21/117). Collectively, these data provide a comprehensive list of putative legumain substrates in native conditions, shed light on cleavage preferences for legumain, and highlight divergent proteolysis in the absence of legumain.

### Putative legumain substrates exhibit extra-lysosomal localisation, potentiating legumain proteolytic activity in neutral environments

We next investigated the 119 asparagine-specific cleavage events (corresponding to 110 proteins) that were enriched in WT spleens, which most likely to result from direct cleavage by legumain (**Fig. 4c**). We plotted the 20 most differential neo-N-termini as a heatmap and observed clear reproducibility across biological replicates (**Supp. Fig. 13f**). Ninety-three (78%) of the mouse cleavage sites exhibit a conserved asparagine at the P1 position of the corresponding human proteins, indicating conservation across species (**Supplementary Table 23)**. When examining the localisation of the putative substrates, only six are catalogued in UniProt as having endo-lysosomal localisation (**Fig. 5a; Supplementary Table 23**). Instead, the majority (76%) are known to be localised to the nucleus and/or cytoplasm. Indeed, STRING (v.11.5) analysis of these proteins indicated a high proportion with GO terms associated with the nucleus (GO:0005634, strength = 0.35, Benj. Hoch. FDR = 6.52e^-11^) (**Fig. 5b**).

We next aimed to validate three putative legumain substrates using an *in vitro* cleavage assay: lysosomal α-mannosidase (MAN2B1; cleaved at N^424^ ↓ V^425^), which was also identified in the RAW264.7 dataset, lamina-associated polypeptide 2 (TMPO; cleaved at N^58^ ↓ S^59^) and tyrosyl tRNA-synthetase 1 (YARS1; cleaved at N^357^ ↓ S^358^) (**Table 1**; **Fig. 6a, 6d, 6g**). We tested the ability of legumain to directly mediate these cleavages by incubating recombinant human proteins in the presence or absence of human legumain. Consistent with our *in vivo* identified N-terminal cleavage events, legumain treatment resulted in notable alterations in gel mobility of the putative substrates (**Fig. 6b, 6e, 6h**). We used dimethylation-based N-terminomics to identify these *in vitro* cleavage sites. These results support the cleavage of TMPO (N^58^ ↓ S^59^) (**Fig. 6f, Supplementary Fig. 14a**) and YARS1 (N^357^ ↓ S^358^) (**Fig. 6i, Supplementary Fig. 14b**) at the residues observed *in vivo*. MAN2B1, on the other hand, was cleaved at alternative sites to those observed *in vivo* (N^630^ ↓ Q^631^, N^665^ ↓ Q^666^, and N^748^ ↓ G^749^; **Fig. 6c; Supplementary Table 12**). Overall, our data demonstrate that legumain possesses the capacity to process these recombinant proteins *in vitro* and validates the use of FAIMS-enabled N-terminomics for accurate identification of novel protease substrates.

**Fig. 6.**
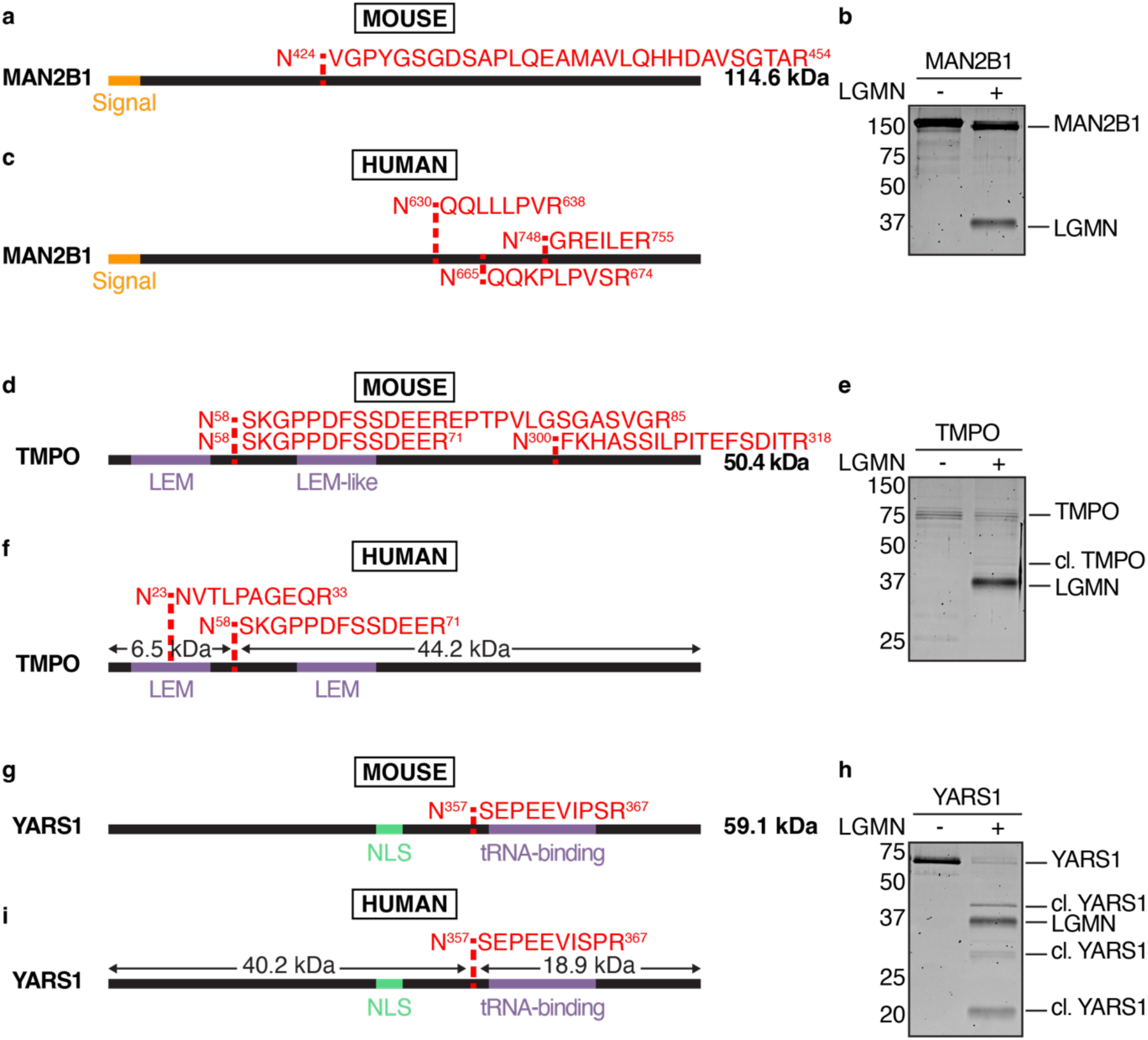
Validating putative legumain substrates identified by FAIMS-enabled N-terminomics. a, d, g. Schematic of murine lysosomal α-mannosidase (MAN2B1) cleavage at N424 ↓ V425 (**a**), lamina-associated polypeptide 2 (TMPO) cleavage at N58 ↓ S59 (**d**), and tyrosyl-tRNA synthetase 1 (YARS1) cleavage at N357 ↓ S358 (**g**) as identified by FAIMS-enabled N-terminomics. Cleavage sites are shown in red, signal peptide in orange, key domains in purple, and nuclear localisation signal (NLS) in green. **b, e, h.** SYPRO Ruby-stained 15% SDS-PAGE gel of recombinant human proteins incubated with (+) or without (-) activated recombinant legumain (*Lgmn*) at pH 5.5 for 5 hours (1:1 mass ratio). Image is representative of 3 independent experiments with similar results. **c, f, i.** Schematic representation of human lysosomal α-mannosidase (MAN2B1) (**c**), lamina-associated polypeptide 2 (TMPO) (**f**), and tyrosyl-tRNA synthetase 1 (YARS1) (**i**) asparaginyl cleavages as identified by N-terminomics analysis of recombinant protein cleavage assays. All dimethylated N-termini containing asparaginyl cleavages detected by LC-MS/MS analysis are shown in red, signal peptide in orange, key domains in purple, and nuclear localisation signal (NLS) in green. Predicted size of resulting cleavage products are shown. For the full dataset, see Supplementary Table 12.

**Table 1.**
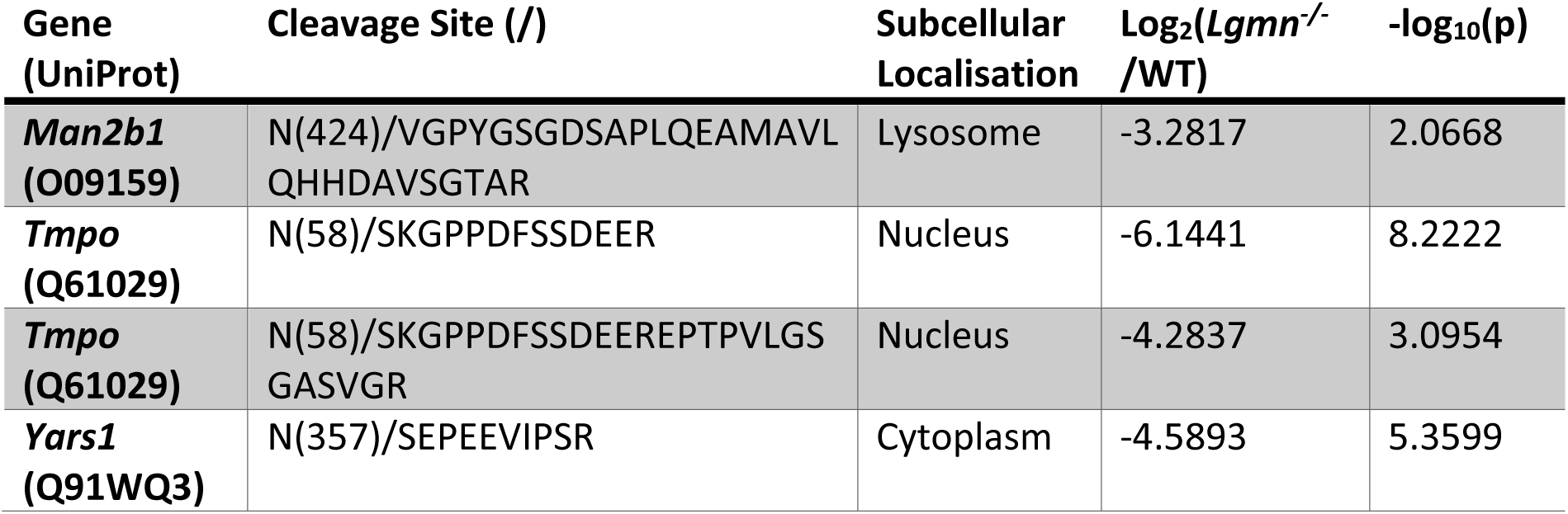
Selected legumain substrates identified in wild-type and legumain-deficient naïve mouse spleens by FAIMS-enabled N-terminomics.

## Discussion

Protease substrate identification is crucial for understanding the complete proteolytic potential of proteases. Although conventional N-terminomics workflows provide sufficient data for *in vivo* substrate identification^19^, they are hampered by difficult N-termini enrichment methods, which often compromise protein abundance information. Here, we developed and employed a novel N-terminomics workflow using FAIMS fractionation^40,41^ for the sensitive identification of both protein abundance changes and cleavage events in RAW264.7 murine macrophages following legumain inhibition. We validated the effectiveness of FAIMS-enabled N-terminomics analysis by benchmarking our approach against unfractionated cell lysates, affirming robust increases in peptide identification and total proteome coverage (**Fig. 1-2**). Additionally, our data highlight the ability of FAIMS to unveil peptides and neo-N-termini unique to each fraction (**Fig. 1c-d**). These data, coupled with the reduced sample handling for LC-MS/MS analysis, validate FAIMS-enabled N-terminomics as an effective strategy for deep proteome and N-terminome analyses, including identification of protein abundance changes and cleavage events.

Using FAIMS-enabled N-terminomics, we investigated global proteome changes and physiological cleavage events in WT and *Lgmn^-/-^* naïve spleen lysates, aiming to identify legumain substrates (**Fig. 3-5**). We chose to analyse murine spleens as legumain has been reported to be highly expressed within this tissue, yet its proteolytic impact has been largely unexplored^54, 77, 79^. We identified a total of 6,366 proteins, revealing global changes to several lysosomal cathepsins in the absence of legumain (**Fig. 3**). As with previous studies, this may suggest compensation for loss of lysosomal hydrolase activity in legumain-deficient samples, which may contribute to lysosomal storage disorders^56^. Martinez-Fábregas and colleagues observed increased expression of lysosomal proteases and hydrolases in legumain-deficient kidneys (including cathepsin A, B, C, L, and X/Z), which is likely driven by STAT3 activation as a response to oxidative stress^57^. In our study, we observed increased cathepsin B, L, E, and K expression in *Lgmn^-/-^* spleens, which may signify tissue-specific responses. The 116 neo-N-termini enriched in *Lgmn^-/-^* spleens (**Fig. 4b**) likely reflect these altered proteolytic networks upon loss of legumain.

Considering cathepsins are involved in various immune processes such as toll-like receptor processing^80^, inflammasome activation^81^, and MHC-II invariant chain processing for antigen presentation^82^, it is unsurprising we see an enrichment of proteins involved in immune-related pathways in *Lgmn^-/-^* spleens (**Fig. 3h**). We also observed upregulation of several neutrophil-associated proteins, including neutrophil elastase and myeloperoxidase (**Fig. 3b-g**). Previous studies have indicated increased populations of Gr-1^+^/Mac-1^+^ cells in *Lgmn^-/-^* spleens, which may be the result of extramedullary haematopoiesis and splenomegaly^79^. While we expect that the increased total abundance of neutrophil proteins is the result of increased neutrophil numbers, it still remains to be investigated whether legumain also mediates cell-intrinsic effects within neutrophils.

To investigate legumain-dependent alterations of the physiological N-terminome, we used N- terminal dimethylation to chemically label native and neo-N-terminal peptides, providing a chemical marker of proteolysis. Making use of the enhanced proteome depth afforded by FAIMS and recent advances in bioinformatics tools (e.g., MSFragger), we demonstrated that cleavage sites could be identified without direct enrichment of N-termini. We observed 119 cleavage events enriched in WT tissue that correspond to cleavage after asparagine residues, representing potential legumain substrates (**Supplementary Table 23**). Whilst other studies have interrogated the degradome of legumain^73, 74, 83^, these studies were performed *in vitro* against a denatured proteome at low pH. Our study is the first to systematically profile the legumain substrate repertoire under native physiological conditions within tissue. While 93 of the cleavage sites are conserved between mouse and human proteins, it will be critical to consider the 26 divergent sites when translating legumain function in mouse models to human pathophysiology.

Intriguingly, only six of the identified cleavage events occurred within the endo-lysosomal system. This may reflect the rapid turnover of substrates in the lysosome, where legumain activity is optimal due to the low pH environment. This may also be the reason that we did not observe enrichment of cleavages after aspartic acid residues, which requires a low pH. The resulting fragments may also be short-lived due to secondary cleavages by other lysosomal proteases. The enrichment of asparagine in the P3 position of the non-P1 asparagine cleavages (**Supplementary Fig. 13e**) may hint at dipeptidyl aminopeptidase activity following legumain cleavage. Cathepsin B has recently been characterized as a dipeptidyl carboxypeptidase with a preference to cleave substrates bearing asparagine in the P2’ position^84^. We hypothesise that secondary cleavage by cathepsin B may obscure detection of legumain substrates in the lysosome, although analysis of the C-terminome would be required to confirm this.

We identified 84 substrates (76%) with known localization to the nucleus or cytoplasm, suggesting a much broader substrate repertoire in these compartments than previously appreciated (**Fig. 5a**). From our data, we cannot confirm the location at which these proteins are cleaved; it is possible that the nuclear/cytoplasmic proteins are cleaved within lysosomes. The lack of identification of known lysosomal proteins, however, provides support that the cleavages occur extra-lysosomally. Legumain localises to the nucleus in the setting of colorectal cancer^52^ and can cleave the nuclear protein FOXP3 in regulatory T-cells to inhibit T- cell differentiation^72^. Cytoplasmic legumain is often associated with neurodegenerative phenotypes, where it cleaves tau^58^, α-synuclein^85^, and SET^86^ to promote neurofibrillary tangles, plaque formation, and cognitive impairment. In the context of Alzheimer’s disease, legumain phosphorylation at S^2^^26^ by SRPK2 led to accumulation of cytoplasmic legumain, promoting cleavage of tau, APP, and SRPK2 itself^87^.

Although our data provide additional support that legumain is able to cleave substrates in neutral environments, how this occurs is still poorly understood. *In vitro*, cystatin E, an endogenous inhibitor of legumain, can bind and stabilise the active conformer at neutral pH by mimicking the C-terminal propeptide^88^. Extracellular legumain can bind to αvβ3 integrin through its RGD motif, which stabilises its activity at neutral pH. Phosphorylation at S^2^^26^ may also function to stabilise legumain in the cytoplasm or nucleus^87^. In any case, we predict that legumain activity is lower in the nucleus or cytoplasm than in the lysosome. This slower cleavage may mediate limited proteolytic events, leading to longer lived products than in the degradative environment of the lysosome. We predict that many of these long-lasting cleavage products will have concerted effects on protein function and cellular processes.

Amongst the putative legumain substrates that we identified, we validated cleavage of human cathepsin S at N^1^^12^ ↓ R^113^ (N^120^ ↓ R^121^ in mouse), TMPO at N^58^ ↓ S^59^, and YARS1 at N^357^ ↓ S^358^ (**Fig. 2g-i**, **Fig. 6**). Cathepsin S is a lysosomal cysteine protease that is involved in antigen presentation^89^ and contributes to diseases such as colitis^90^, inflammation^91^, and oral cancer^92^. Various studies have previously reported legumain activity to be essential in the processing of several lysosomal cathepsins, including cathepsin B, L, D, and H^56, 57, 93^. Our data suggest that legumain activity may also regulate the maturation of cathepsin S (**Fig. 2**). Interestingly, cleavage of mouse cathepsin S at N^1^^20^ ↓ R^121^ was detected in RAW264.7 cells, but not spleen. In both datasets, cleavages at R^121^ ↓ T^122^ and T^122^ ↓ L^123^ were also detected. The latter is the predicted cleavage site for removal of the cathepsin S pro-peptide to facilitate its activation.

These results suggest that in some contexts, legumain may directly mediate cathepsin S activation. As N^120^RT is a predicted site for *N*-linked glycosylation on mouse cathepsin S, we hypothesise that differential modification of this residue may dictate whether it can be cleaved by legumain. In human cathepsin S, however, the glycosylation site (N^104^IT) is distinct from the legumain cleavage site (N^112^ ↓ R^113^), suggesting differences in the interplay of these proteases across species.

TMPO is responsible for maintenance of the nuclear envelope and association with chromatin^94^. We observed degradation of TMPO in the presence of legumain *in vitro* (**Fig. 6e**). We hypothesise that in the neutral environment of the nucleus, legumain would cleave TMPO in a more controlled manner. Caspase-3 and -6 have been implicated in processing TMPO^95,96^, resulting in release of chromatin from the nuclear envelope for degradation during early stages of apoptosis^97^. Considering the identified N^58^ ↓ S^59^ cleavage is located between a LEM domain and the chromatin binding region, it is plausible that similar results occur in the presence of legumain. The relationship between legumain and TMPO therefore requires further investigation.

YARS1 is a ligase that catalyses the attachment of tyrosine to tRNA molecules^98^. The legumain cleavage site on YARS1 (N^3^^57^ ↓ S^3^^58^) is located between its tRNA binding domain and nuclear localisation signal (NLS) (**Fig. 6g-i**). Considering both domains are required for Tyr-tRNA binding during protein synthesis^99^, we hypothesise that legumain cleavage may lead to reduced tyrosine-tRNA ligase activity. The NLS may also become unmasked following legumain cleavage, thereby increasing the nuclear activities of YARS1^98^. Aminoacyl-tRNA synthetases are also known to be processed into fragments with cytokine potential. In the case of YARS1, matrix metalloproteinase (MMP)-mediated cleavage at S^386^ ↓ L^387^L and G^405^ ↓ L^406^ can enhance TLR2 signalling, TNF-α secretion from macrophages, and amplify monocyte/macrophage chemotaxis^100^. YARS1 processing typically separates the N-terminal Rossman fold and C- terminal EMAPII domain (365-528 aa), yielding an N-terminal fragment (mini-TyrRS) known to promote endothelial cell migration and angiogenesis through transactivation of vascular endothelial growth factor receptor-2 (VEGFR2)^101,102^. N^357^ ↓ S^358^ was the only cleavage product identified in our spleen dataset, and it was 25-fold enriched in WT tissue compared to *Lgmn^-/-^*. Interestingly, this site was also enriched in inflamed skin sections from *Mmp2^-/-^* mice when compared to wild-type^103^. Considering legumain can cleave and activate pro-MMP-2^70^, legumain may be upregulated in MMP2-deficient mice to compensate its loss, and consequently, YARS1 is more processed. It is plausible that legumain cleavage at this site can mediate cytokine-like effects, but further study is required to test this.

Our *in vitro* cleavage assay on MAN2B1 demonstrated a slight shift in mobility in the presence of legumain (**Fig. 6b**). As this is not indicative of N^424^ cleavage, we hypothesise MAN2B1 may instead be trimmed in this assay (**Fig. 6a-b**). Intriguingly, human MAN2B1 is known to undergo post-translational processing into five smaller polypeptides (A-E). Cleavage of G^429^ ↓ S^430^ separates the A and B subunits^104^. This site, along with N^424^ ↓ V^425^ and V^425^ ↓ G^426^ were significantly enriched in wild-type spleens and may suggest increased MAN2B1 processing in the presence of legumain with redundancy in the specific cleavage site (**Supplementary Table 17**). Considering processing into the ABC polypeptide occurs prior to separation of A and B subunits, it is plausible that legumain access may be blocked in our *in vitro* assay and we instead observe cleavage at different sites (**Fig. 6c**). The presence of these processing events in the murine proteome however, are yet to be elucidated and require further investigation.

In summary, we have validated the use of FAIMS-enabled N-terminomics analysis for the robust and streamlined detection of protein abundance changes and cleavage events in wild- type and legumain-deficient mouse spleens. We identified a range of altered proteins including lysosomal cathepsins and neutrophil-associated proteins. Moreover, we provided the first comprehensive list of physiological legumain substrates identified using a systematic and unbiased approach, revealing novel insights into the proteolytic potential of legumain, especially outside of its lysosomal functions. These studies will assist in the delineation of the complete function of legumain in the cell and support efforts to develop legumain-targeted therapeutics for cancer and neurodegenerative diseases.

## Methods

### Cell culture

RAW264.7 cells (mouse monocyte/macrophage) were cultured in Dulbecco’s Modified Eagle Medium (DMEM, high glucose, Thermo) supplemented with 10% Foetal Bovine Serum (FBS, CellSera) and 1% antibiotics (100 U/mL penicillin-streptomycin, Thermo) at 37 °C with 5% CO2. Cells were passaged 1:10 once reaching 80-90% confluence using a cell scraper.

### Mice

*Lgmn*^-/-^ C57BL/6N were a gift from Thomas Reinheckel^105^. Mice were bred in the laboratory of Brian Schmidt and Nigel Bunnett at New York University and studies were approved by the NYU Institutional Animal Care and Use Committee. Splenic tissues were harvested from 8- week-old healthy male mice (WT and *Lgmn^-/-^*), snap frozen and stored at -80 °C.

### Inhibition of legumain and assessment of legumain activity

The legumain-specific inhibitor SD-134^75^ was used to inhibit legumain in RAW264.7 cells. Cells (2x10^6^) were seeded in a 6-well plate, followed by addition of vehicle or SD-134 (10 µM added from a 10 mM stock; 0.1% final DMSO concentration). After 16 hours, cells were harvested and used for MS analysis as below. Alternatively, the activity-based probe LE28^77^ was used to assess residual legumain activity and inhibitor efficacy. Cells were lysed in citrate buffer (50 mM citrate (Thermo, pH 5.5), 0.5% CHAPS (Sigma), 0.1% Triton X-100, 4 mM DTT (Sigma)), and solids were cleared by centrifugation (21,000 x *g*, 5 minutes, 4 °C). Protein concentrations in the resulting supernatant were determined by bicinchoninic acid (BCA) assay according to manufacturer’s instructions (Pierce). Total protein (100 µg) was diluted in 20 µL citrate lysis buffer and LE28 (1 µM) was added from a stock of 100 µM (1% final DMSO concentration). After 30 minutes at 37 °C, the reaction was quenched by addition of 5x sample buffer (50% glycerol (Sigma), 250 mM Tris-Cl (Sigma), pH 6.8, 10% SDS (VWR LifeSciences), 0.04% bromophenol blue (Sigma), 6.25% beta-mercaptoethanol (Sigma), diluted to 1x final). Samples were boiled at 95 °C for 5 minutes, and proteins were resolved on a homemade 15% SDS-PAGE gel. To detect LE28-labelled species, gels were scanned with a Cy5 filter set on a Typhoon flatbed laser scanner (GE Healthcare). Spleens were similarly analysed with LE28 to confirm loss of activity in *Lgmn^-/-^* tissue, except lysis in citrate buffer was facilitated by sonication.

### ImmunobloDng

Proteins were transferred to nitrocellulose membranes using the Trans-Blot Turbo Transfer system (Biorad) and incubated with the indicated primary antibody overnight at 4 °C. Blots were washed in PBS containing 0.1% Tween-20 (PBST; Sigma) three times before incubation with the secondary antibody for one hour at room temperature and three washes with PBST.

A final wash in PBS was performed prior to detection. Horseradish peroxidase (HRP) conjugated antibodies were detected with Clarity ECL Substrate (Biorad) on a ChemiDoc (Biorad). Fluorescent-conjugated antibodies were visualised using the Typhoon 5 IRlong channel. Ponceau S stain was used to evaluate loading and transfer efficacy. All antibodies were diluted in 1:1 Intercept blocking buffer (LI-COR) and PBST. Bands were quantified by densitometry using ImageJ (Fiji) with background subtraction. Antibodies used in this study included; goat anti-mouse legumain (1:1,000, R&D AF2058), goat anti-mouse elastase 2A (1:1,000, R&D AF4517), rabbit anti-β-actin (1:10,000, Life Technologies, MA5-15739), donkey anti-goat IgG HRP-conjugated (1/10,000, Novex Life Technologies, A15999), goat anti-rabbit IgG IR800-conjugated (1:10,000, Li-cor, 926-32213).

### In vitro recombinant protein cleavage assay

To assess the direct cleavage of various proteins by legumain *in vitro,* we incubated recombinant proteins (Supplementary Table 24) with activated recombinant human legumain (1.5 µg/µL stock, gifted by Hans Brandstetter) in acetate buffer (50 mM sodium acetate (ChemSupply), 100 mM sodium chloride (EMSURE), pH 5.5) and incubated at 37 °C for 5 hours. Samples in the absence of legumain were used as a negative control. The reactions were quenched by addition of 5x sample buffer (1x final concentration) prior to analysis by SDS-PAGE. Band visualisation was achieved using SYPRO™ Ruby Gel Stain (Invitrogen). Briefly, gels were fixed in 50% methanol, 7% acetic acid twice for 30 minutes at room temperature with shaking followed by overnight staining with SYPRO™ Ruby Gel Stain. Gels were washed in 10% methanol, 7% acetic acid for 30 minutes at room temperature followed by two 5-minute washes in Mill-Q water for imaging on the ChemiDoc (BioRad). Alternatively, following a 5-hour incubation at 37 °C, samples were quenched with 4% SDS, 250 mM Tris-HCl (pH 6.8) and prepared for N-terminomic analysis.

### Protein N-termini dimethylation and proteome preparation

RAW264.7 cells were treated with DMSO or 10 µM SD-134 as above and harvested (n = 4/group). Splenic tissue was harvested from wild-type C57BL/6 and *Lgmn^-/-^* mice and stored at -80 °C (n = 4/group). Cells and tissues were lysed by sonication in 4% SDS, 50 mM HEPES (pH 7.5, Sigma) containing Roche cOmplete, EDTA-free protease inhibitor (Sigma). After boiling for 10 minutes, lysates were cleared by centrifugation (21,000 x *g*, 5 minutes, 4 °C) and total protein (100 µg) was diluted in 100 µL buffer according to BCA analysis. Recombinant proteins from the *in vitro* cleavage assay were prepared as described above.

Proteins were reduced with 20 mM DTT (80 °C, 10 minutes, 500 rpm) and alkylated with 50 mM iodoacetamide (37 °C, 30 minutes, 500 rpm) in the dark followed by quenching with 50 mM DTT (37 °C, 20 minutes, 500 rpm). Paramagnetic beads (Sera-Mag SpeedBeads 45152105050250 and 65152105050250, GE Healthcare) were prepared by mixing in a 1:1 ratio and washing three times in Milli-Q water before adjusting to a final concentration of 50 µg/µL in Milli-Q water as previously outlined^106^. Conditioned SP3 beads were added to samples (2 mg of SP3 beads, final protein:SP3 bead ratio of 1:20) and protein aggregation was initiated with the addition of ethanol (80% final concentration). Samples were then gently shaken (25 °C, 1,000 rpm) for 20 minutes prior to washing three times with 500 µL of 80% ethanol using a magnetic rack and resuspending in 90 µL of 6 M guanidine hydrochloride, 100 mM HEPES (pH 7.5). Proteins were dimethylated by adding 30 mM formaldehyde (Sigma) and 30 mM sodium cyanoborohydride (Sigma) and shaking (37 °C, 1,000 rpm) for 1 hour. This was repeated once more with an additional 30 mM of formaldehyde and 30 mM sodium cyanoborohydride before labelling was quenched by adding 25 µL 4 M Tris-base (pH 6.8) and shaking (37 °C, 1,000 rpm) for 1 hour. Excess formaldehyde and sodium cyanoborohydride were removed from samples using SP3 clean up (1 mg SP3 beads; final protein:SP3 bead ratio of 1:30) and proteins were precipitated with ethanol (80% final concentration). Samples were gently shaken (25 °C, 1,000 rpm) for 20 minutes and then washed three times with 500 µL of 80% ethanol using a magnetic rack. SP3 beads were then resuspended in 100 μL of 200 mM HEPES (pH 7.5) and digested overnight at 37 °C with Solu-trypsin (3 µg solu-trypsin, Sigma, trypsin:protein ratio 1:33). The resulting peptide mixtures were collected using a magnetic rack, acidified with Buffer A* (0.1% trifluoroacetic acid, 2% acetonitrile) and desalted using C18 StageTips (Empore™, 3M) with the addition of Oligo-R3 resin reverse phase material (Thermo) as previously described^107,108^. Samples were dried using a speedvac and stored at - 20 °C until analysis.

### Online fractionation by high-field asymmetric waveform ion mobility spectrometry (FAIMS) and mass spectrometry (MS) analysis

Proteome samples were re-suspended in Buffer A* and separated using a two-column chromatography setup composed of a PepMap100 C18 20-mm by 75-μm trap and a PepMap C18 500-mm by 75-μm analytical column (Thermo Fisher Scientific) on a Dionex Ultimate 3000 UPLC (Thermo Fisher Scientific). Samples were concentrated onto the trap column at 5 μL/min for 5 min with Buffer A (0.1% formic acid, 2% DMSO) and then infused into an Orbitrap 480™ mass spectrometer (Thermo Fisher Scientific) equipped with a FAIMS Pro interface at 300 nL/minute. For each sample/FAIMS fraction ∼2 µg of peptide mixtures were separated using 125-minute analytical runs undertaken by altering the buffer composition from 3% Buffer B (0.1% formic acid, 77.9% acetonitrile, 2% DMSO) to 23% B over 95 min, then from 23% B to 40% B over 10 min, then from 40% B to 80% B over 5 min. The composition was held at 80% B for 5 min, and then dropped to 2% B over 0.1 min before being held at 2% B for another 9.9 min. For each sample, six individual LC-MS runs were collected with the Orbitrap 480™ Mass Spectrometer operated using different FAIMS compensational voltages (CV) of either -35, -45, -55, -65, -75 or -85. For each FAIMS fraction, data-dependent acquisition was undertaken with a single Orbitrap MS scan (300-2,000 m/z, a resolution of 120k with the Automated Gain Control (AGC) set to a maximum of 300%) collected every 3 seconds followed by Orbitrap MS/MS HCD scans of precursors (Normalized collision energy of 30%, maximal injection time of 50 ms, a resolution of 30k and a AGC of 250%). Non-FAIMS analysis was undertaken using the same LC-MS parameters as outlined above on the same biological samples used for FAIMS analysis.

Dimethylated and trypsin digested *in vitro* cleavage assay samples were re-suspended in Buffer A* and separated on a two-column chromatography setup composed of a PepMap100 C18 20-mm by 75-μm trap and a PepMap C18 500-mm by 75-μm analytical column (Thermo Fisher Scientific) on a Dionex Ultimate 3000 UPLC (Thermo Fisher Scientific) coupled to a Q Exactive Plus Orbitrap mass spectrometer (Thermo Fisher Scientific). Each sample (3 µg) of peptide mixtures were injected into the mass spectrometer at 300 nL/min and separated using 65-minute analytical runs undertaken by altering the buffer composition from 2% Buffer B to 23% B over 35 min, then from 23% B to 40% B over 10 min, then from 40% B to 80% B over 5 min. The composition was held at 80% B for 5 min, and then dropped to 2% B over 0.1 min before being held at 2% B for another 9.9 min. Data-dependent acquisition was undertaken with a single Orbitrap MS scan (375-2,000 m/z, a resolution of 70k with the Automated Gain Control (AGC) target set to 3 x 10^6^ and maximal injection time of 50 ms) followed by targeted analysis with parallel reaction monitoring (PRM) for sensitive identification of specific peptides (Stepped HCD Normalized collision energy of 30%, 35%, and 40%, resolution of 35k with a AGC target set to 3 x 10^5^ and maximal injection time of 150 ms)^109^. Inclusion criteria of the peptides of interest were specified based on the double, triple, or quadruple-charged states of the human protein sequences (Supplementary Table 25).

### Quantitative proteomics and N-terminomics data analysis

RAW264.7 cell lysate and murine spleen lysate data files were processed and searched using MSFragger (Fragpipe v.18.0)^76^ against the unreviewed mouse proteome (*Mus musculus,* UniProt Accession: UP000000589), supplemented with common contaminants, and a reverse decoy database. Identification and label free quantification (LFQ) was undertaken allowing for cysteine carbamidomethylation as a fixed modification (+57.0215 Da) as well as variable modifications of lysine dimethylation (+28.0313 Da), methionine oxidation (+15.9949 Da), N- terminal acetylation (+42.0106 Da), N-terminal cyclisation (-17.0265/-18.0106 Da), N-terminal dimethylation (+28.0313 Da), and N-terminal lysine dimethylation (+56.0626 Da). Cleavage specificity was set as “TrypsinR” (Arg-C), allowing a maximum of 2 missed cleavages. A precursor mass tolerance of 20 ppm and isotopic error of 3 Da were also included. The resulting outputs were further processed in Perseus (v.1.6.0.7)^110^, removing reverse decoy matches and protein contaminants before a log2 transformation was applied. Peptides identified in a minimum of three of four biological replicates in at least one of the groups (DMSO/SD-134 or WT/*Lgmn^-/-^*) were selected and missing values imputed based on a downshifted normal distribution (σ-width = 0.3, σ-downshift = -1.8). Student’s two-sample t-test was applied for statistical comparison between groups. Volcano plots, charts, heatmaps, principal component analyses, upset plots, and Venn diagrams were all created using R (v.4.2.0). Enrichment analysis using Fisher exact tests were undertaken in Perseus and visualisation of proteomic data undertaken in the R statistical environment using the ggplot2 package (v.3.3.6)^111^ in R (v.4.2.0). Pearson correlation and statistical summary analyses were performed in Perseus and standard deviations taken for visualisation.

Recombinant protein cleavage assay data files were searched against a custom human database containing sequences for legumain (UP#Q99538), cathepsin S (UP#P25774), lysosomal α-mannosidase (UP#O00754), lamina-associated polypeptide 2 (UP#P42167), and tyrosyl-tRNA synthetase (UP#P54577) with common contaminants and a reverse decoy database added by MSFragger (Fragpipe v.18.0). Identification and quantification of peptides occurred as described above. *In vitro* cleavage assay spectra were manually assessed and annotated with the Interactive Peptide Spectral Annotator^112^.

### Bioinformatic analysis of protein and peptide data

Data were processed in WebPICS^113^ and TopFINDer^114,115^ for generation of sequence logos using plogo^116^. STRING-dp (v.11.5) was used for protein interaction and pathway analyses (https://string-db.org) with medium confidence (0.400) and FDR stringency (5%).

### Myeloperoxidase (MPO) activity assay

Splenic tissues were sonicated in 50 mM potassium phosphate buffer (pH 6.0) containing 0.5% hexadecyltrimethylammonium bromide (HTAB, Sigma) using the method described above. Total protein was normalised by BCA (7 µg in 7 µL lysis buffer) and aliquoted into a Corning Costar 96-well flat bottom clear plat. Potassium phosphate buffer (50 mM, pH 6.0) containing 0.167 mg/mL O-dianisidine-HCl (Sigma), 0.0005% H_2_O_2_ (Sigma) was added, and absorbance read at 460 nm every 30 seconds for 30 minutes on the Clariostar Omega Plate Reader. Linear values were taken to calculate slopes and graph results.

### Statistical Analysis

Statistical analyses were performed using GraphPad Prism 9 unless otherwise stated. All data are presented as mean ± SEM with significance set at p<0.05. All pairwise comparisons were analysed using a student’s t-test.

### Data Availability

The mass spectrometry proteomics data has been deposited in the Proteome Xchange Consortium via the PRIDE partner repository^117^ with the data set identifiers PXD043136, PXD043124, and PXD043122.

## Acknowledgements

We thank T. Reinheckel for providing access to the legumain-deficient mouse strain and B. Schmidt and N. Bunnett for breeding the mice and providing spleen tissue. We thank E. Dall and H. Brandstetter for the kind gift of recombinant legumain. We thank the Melbourne Mass Spectrometry and Proteomics Facility of The Bio21 Molecular Science and Biotechnology Institute for access to MS instrumentation. L.E.E.-M. was supported by a Grimwade Fellowship funded by the Russell and Mab Grimwade Miegunyah Fund at the University of Melbourne, a DECRA Fellowship from the Australian Research Council (ARC, DE180100418), and a grant from the National Health and Medical Research Council (NHMRC, GNT2011119). N.E.S is supported by an ARC Future Fellowship (FT200100270), an ARC Discovery Project Grant (DP210100362) and a NHMRC Ideas grant (2018980). A.R.Z. was supported by an RTP Scholarship from the Australian Government.

## Author Contributions

A.R.Z. performed experiments, completed the data analysis, and wrote the manuscript. L.E.E.-M. conceived and designed the study, contributed to analysis and interpretation of the study, and edited the manuscript. N.E.S. designed the N-terminomics methods, assisted with analysis, and edited the manuscript. A.D. contributed critical intellectual input and edited the manuscript.

## Competing Interests

All authors have nothing to disclose.

## Notes

### Competing Interest Statement

The authors have declared no competing interest.

