## Supplementary Figures for "FAIMS-enabled N-terminomics analysis reveals novel legumain substrates in murine spleen"

### Supplementary Information

**Supplementary Table 24. Recombinant Proteins used for the *in vitro* cleavage assays.**

| Protein (Gene) | Species | Source | Catalogue Number |
|---|---|---|---|
| Legumain *(Lgmn)* | Human | N/A | Gift from Brandstetter Lab |
| Cathepsin S *(Ctss)* | Human | Mouse myeloma cell line | R&D, 1183-CY-010 |
| Lysosomal alpha-mannosidase *(Man2b1)* | Human | Human embryonic kidney (HEK) cell line | R&D, 986-GH-050 |
| Lamina-associated polypeptide 2 *(Tmpo)* | Human | Human embryonic kidney (HEK293T) cell line | BosterBio, PROTP42167 |
| Tyrosyl-tRNA synthetase 1 *(Yars1)* | Human | *E.coli* | BosterBio, PROTP54577 |

**Supplementary Table 25. Parallel reaction monitoring peptide inclusion list for *in vitro* cleavage assay N-terminomics analysis.**

| Protein (Gene) | Sequence | Charge | m/z |
|---|---|---|---|
| Cathepsin S *(Ctss)* | RILPDSVDWR | 2 | 642.85010 |
| Cathepsin S *(Ctss)* | RILPDSVDWR | 3 | 428.90030 |
| Lysosomal alpha-mannosidase *(Man2b1)* | ANMWFKNLDKLIR | 3 | 578.32940 |
| Lysosomal alpha-mannosidase *(Man2b1)* | ANMWFKNLDKLIR | 4 | 433.99720 |
| Lamina-associated polypeptide 2 *(Tmpo)* | SKGPPDFSSDEER | 2 | 753.85070 |
| Lamina-associated polypeptide 2 *(Tmpo)* | SKGPPDFSSDEER | 3 | 502.90070 |
| Tyrosyl-tRNA synthetase 1 *(Yars1)* | SEPEEVIPSR | 2 | 585.79720 |
| Tyrosyl-tRNA synthetase 1 *(Yars1)* | SEPEEVIPSR | 3 | 390.86510 |

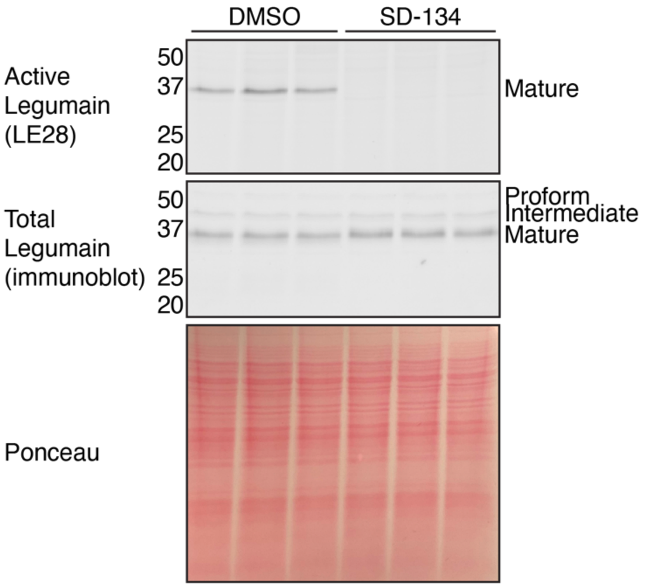


**Supplementary Fig. 1. SD-134 inhibits legumain activity in RAW264.7 cells.** The legumain inhibitor SD-134 (10 µM) was added to RAW264.7 cells 16 hours prior to lysate labelling with the legumain-specific activity-based probe LE28 for 30 minutes (1 µM). In-gel fluorescence of legumain activity was detected by scanning for Cy5 fluorescence on a Typhoon 5 flatbed laser scanner (GE Healthcare). Total legumain was also detected by immunoblot. Ponceau S stain was used as a loading control.

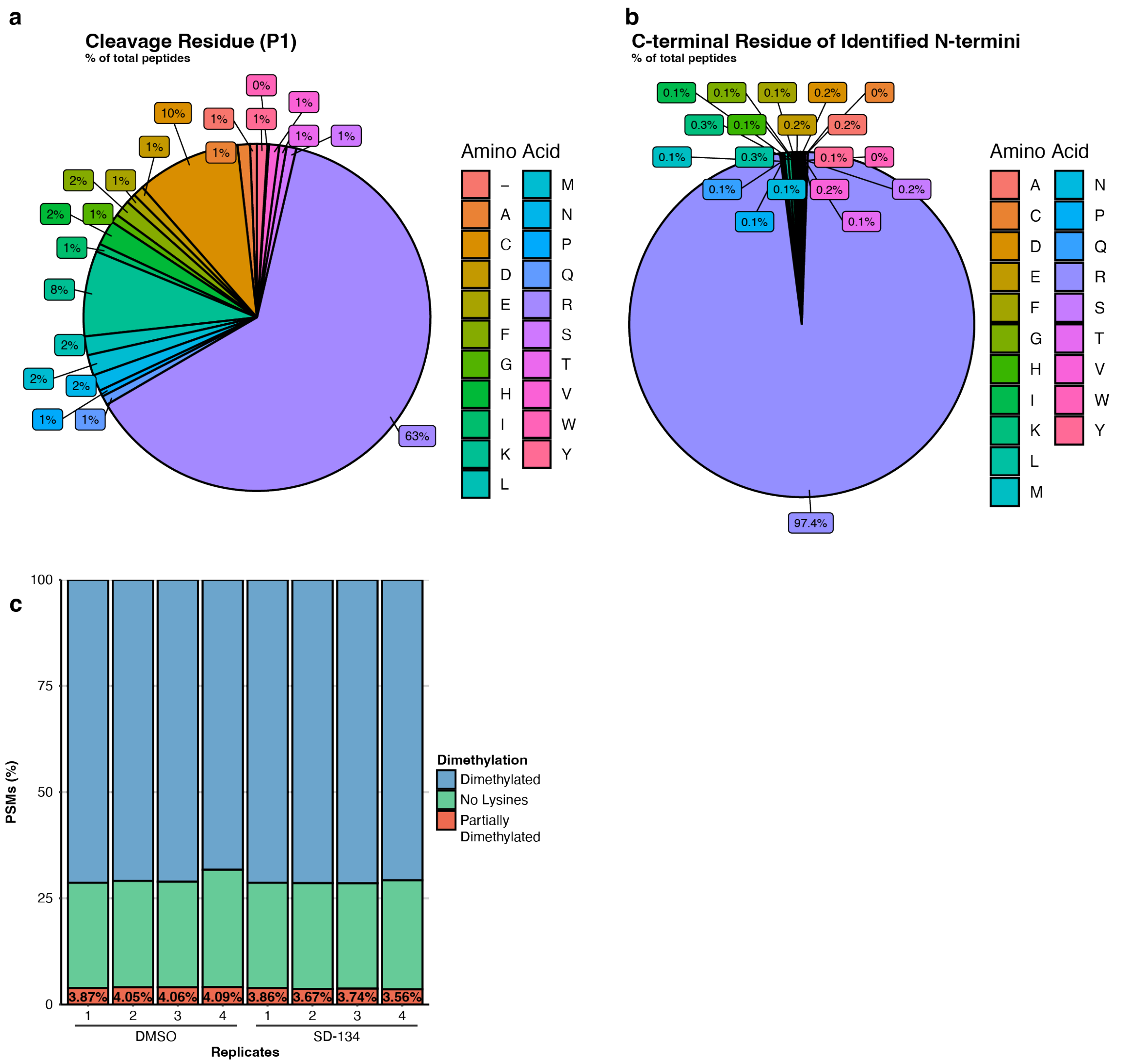


**Supplementary Fig. 2. Dimethylation efficacy of SD-134 (10 µM) and DMSO-treated RAW264.7 cell lysates. a-b.** Cell lysates were denatured, reduced, and alkylated prior to N-terminal dimethylation by formaldehyde. Following LC-MS/MS analysis, amino acid residues prior to the identified peptide/P1 residue (**a**) and at the end of each identified peptide (**b**) were used as measures of dimethylation efficacy. **c.** Dimethylation status of each peptide was also analysed according to whether all lysines were dimethylated (blue), no lysines were present (green), or lysines were partially dimethylated (red) for each biological replicate.

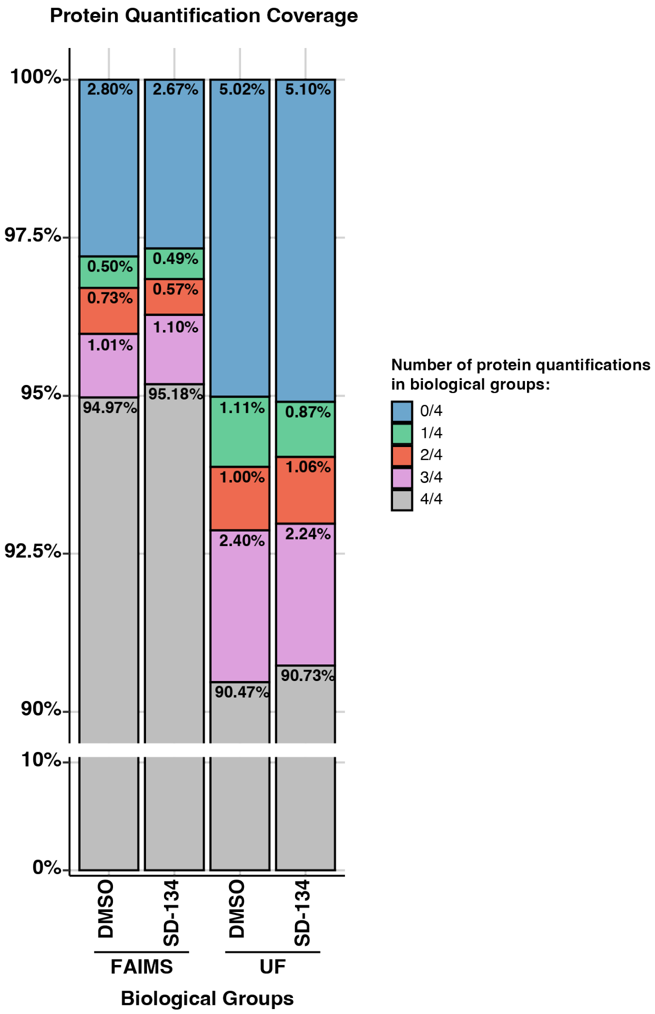


**Supplementary Fig. 3. FAIMS-fractionation of RAW264.7 cell lysates improves proteome coverage of quantified N-termini.** Following LC-MS/MS analysis, data were matched against a murine database in MSFragger for identification and quantification of proteins and N-termini. Quantified N-termini were assessed per biological group in R (v.4.2.0) as to whether they contained protein quantifications in each biological replicate. Zoom shows range from 90-100%.

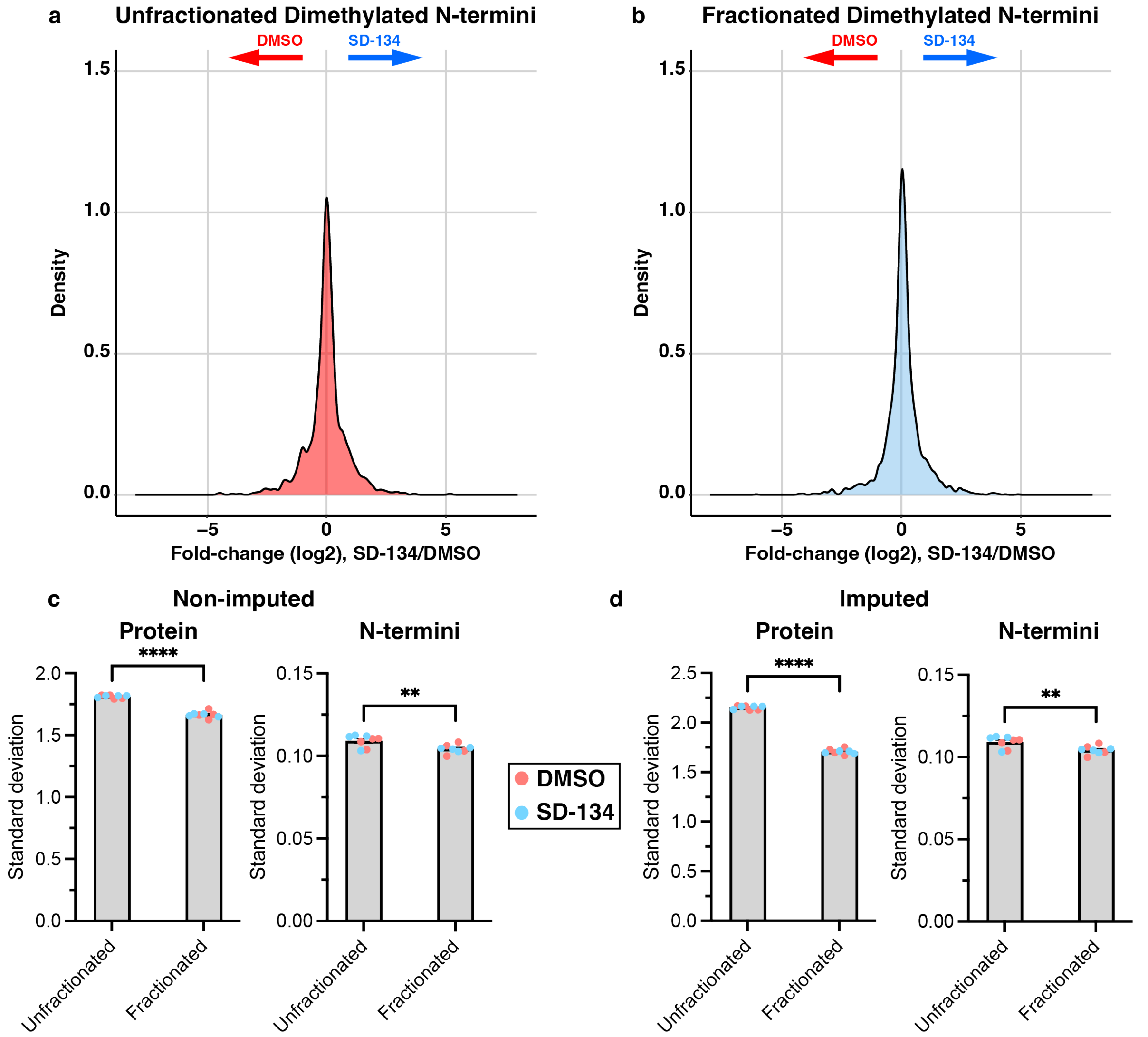


**Supplementary Fig. 4. FAIMS fractionation of RAW264.7 cell lysates enabled tighter distribution of LC-MS/MS data. a-b.** Following LC-MS/MS analysis of FAIMS-fractionated SD-134 (blue) and DMSO (red) treated RAW264.7 cell lysates, data files were processed in Perseus (v.1.6.0.7) to include at least three of four valid values in at least one of the groups. Unfractionated (**a**) and FAIMS-fractionated (**b**) data were visualised as a density plot following a two-way t-test to calculate $\log_2$(SD-134/DMSO) (n = 4/group). **c-d.** Standard deviations of $\log_2$(SD-134/DMSO) were also calculated for each biological replicate on the protein and N-termini level for both non-imputed (**c**) and imputed (**d**) data in Perseus (v.1.6.0.7). Imputed values were based on a normal distribution with σ-width = 0.3 and σ-downshift = -1.8. A student's t-test was used for pairwise comparisons (*$p<0.05$, **$p<0.01$, ***$p<0.001$, ****$p<0.0001$).

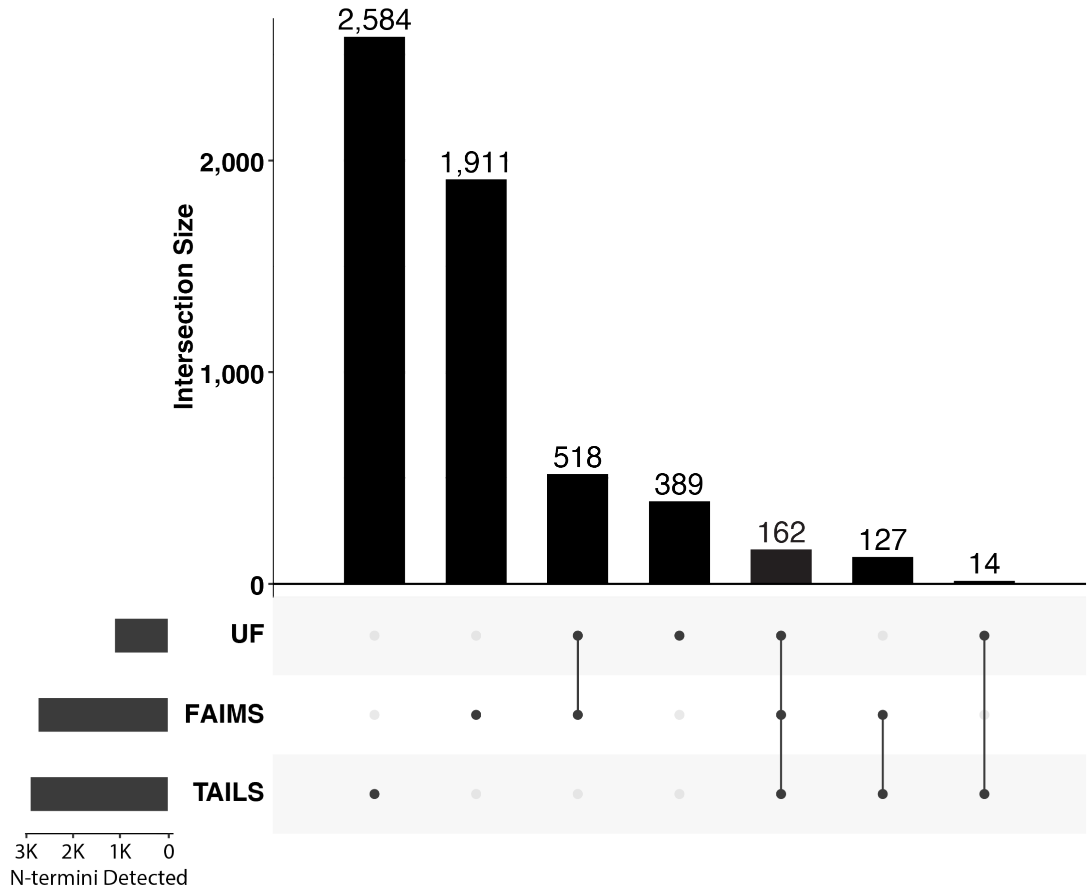


**Supplementary Fig. 5. Comparison of N-termini identifications to a conventional N-terminomics workflow (TAILS).** Overlap of N-termini observed between unfractionated, FAIMS-fractionated, and TAILS (Terminal Amine Isotopic Labelling of Substrates, data obtained from Anderson et al. (2020)) methods. Bottom-left panel shows total N-termini detected in each experiment. UF=unfractionated. The TAILS experiment was performed as followed; RAW264.7 cells were treated with DMSO or 100 µM legumain-specific inhibitor LI-1 (n =4). Following labelling with light and heavy formaldehyde, N-termini were negatively selected for using a dendritic polyglycerol aldehyde polymer for LC-MS/MS analysis on the Orbitrap Fusion Lumos Tribrid mass spectrometer. LC-MS/MS files were searched against the murine proteome database using MaxQuant (v.1.6.0.1) for peptide-spectrum matching.

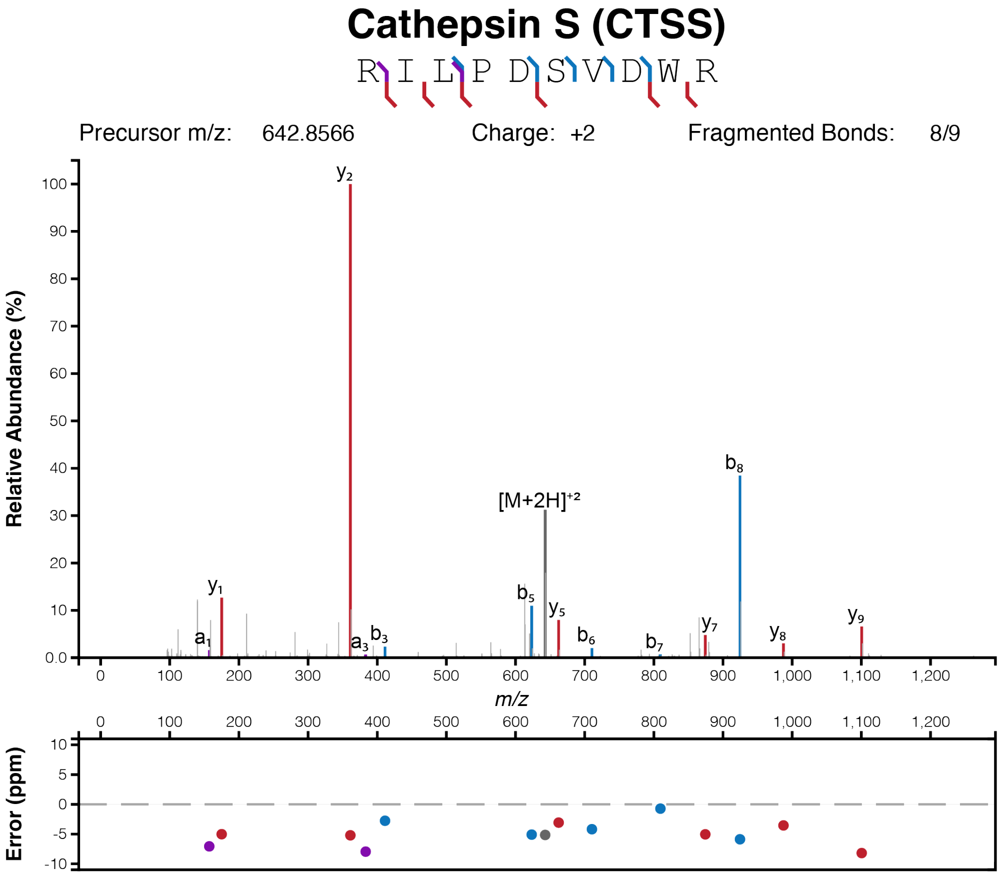


**Supplementary Fig. 6. Legumain directly processes cathepsin S *in vitro* as identified by FAIMS-enabled N-terminomics.** Recombinant proteins were incubated with activated recombinant legumain prior to unfractionated N-terminomics analysis. MS2 analysis revealed dimethylation of the peptide $^{113}$RILPDSVDWR$^{122}$ within legumain-treated cathepsin S samples supporting cleavage at this site. Figures were created using http://www.interactivepeptidespectralannotator.com/PeptideAnnotator.html with a fragment tolerance of ±10 ppm (Schmidt et al. 2021).

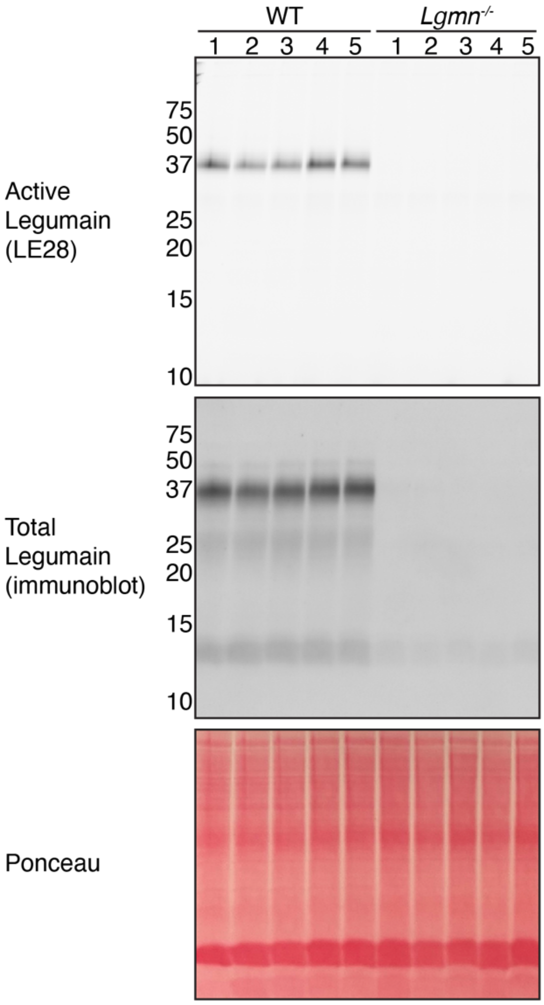


**Supplementary Fig. 7. Legumain activity and expression are lost in legumain-deficient (*Lgmn*$^{-/-}$) naïve mouse spleens.** Spleens were lysed and labelled with the legumain-specific activity-based probe LE28 (1 µM) prior to SDS-PAGE analysis. In-gel fluorescence was imaged with the Cy5 filter of the Typhoon 5 flatbed laser scanner (GE Healthcare). Total legumain was also detected by immunoblot. Ponceau S stain was used as a loading control.

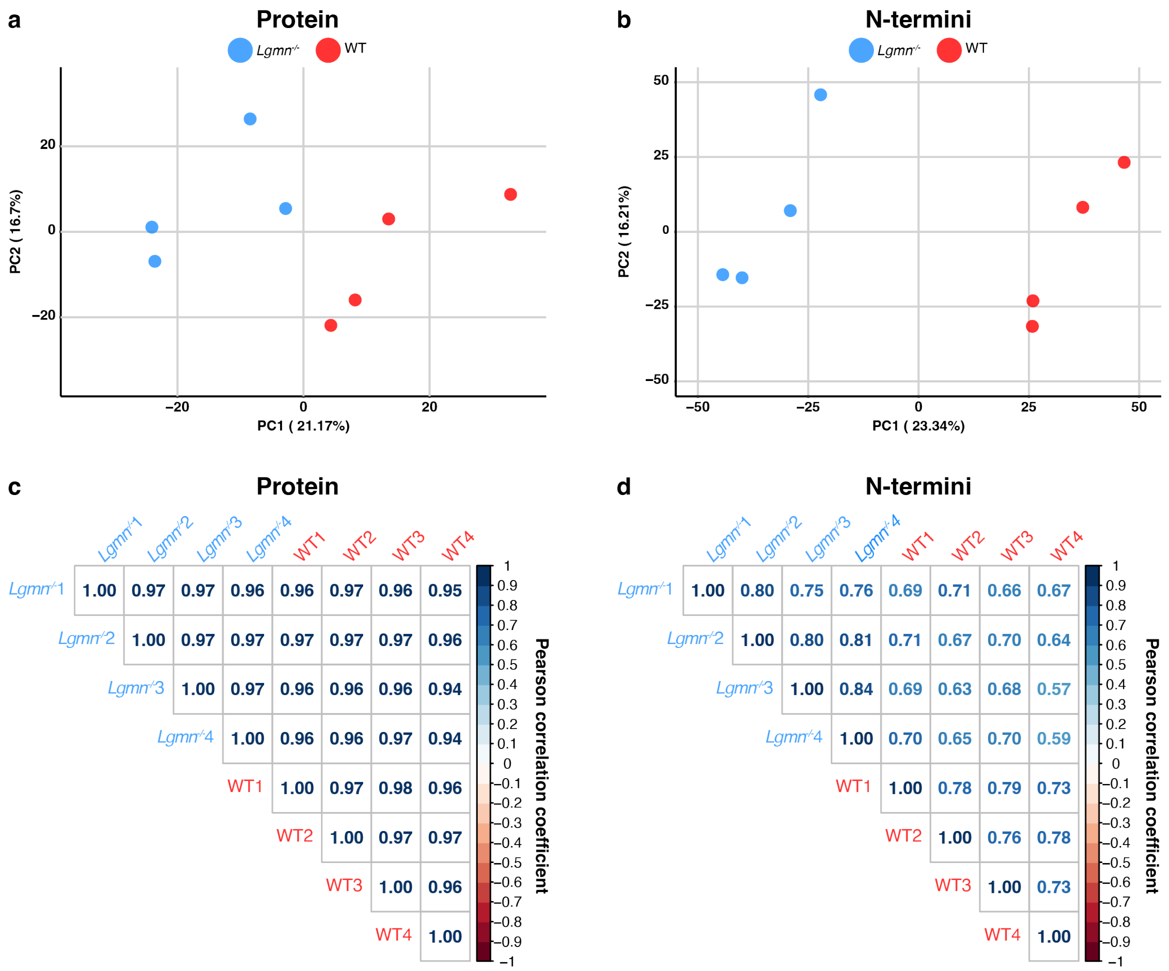


**Supplementary Fig. 8. FAIMS-fractionated spleen lysate LC-MS/MS data demonstrates clustering at biological replicate level. a-b.** Principal component analysis (PCA) of proteins (**a**) and N-termini (**b**) identified using FAIMS-enabled N-terminomics was performed. Wild-stype (WT) data are shown in red and legumain-deficient (*Lgmn*$^{-/-}$) data in blue (n = 4/group). **c-d.** Max label-free quantification values per biological replicate were analysed for Pearson correlation coefficient in Perseus (v.1.6.0.7) and visualised as a correlogram using the corrplot package in the R statistical environment.

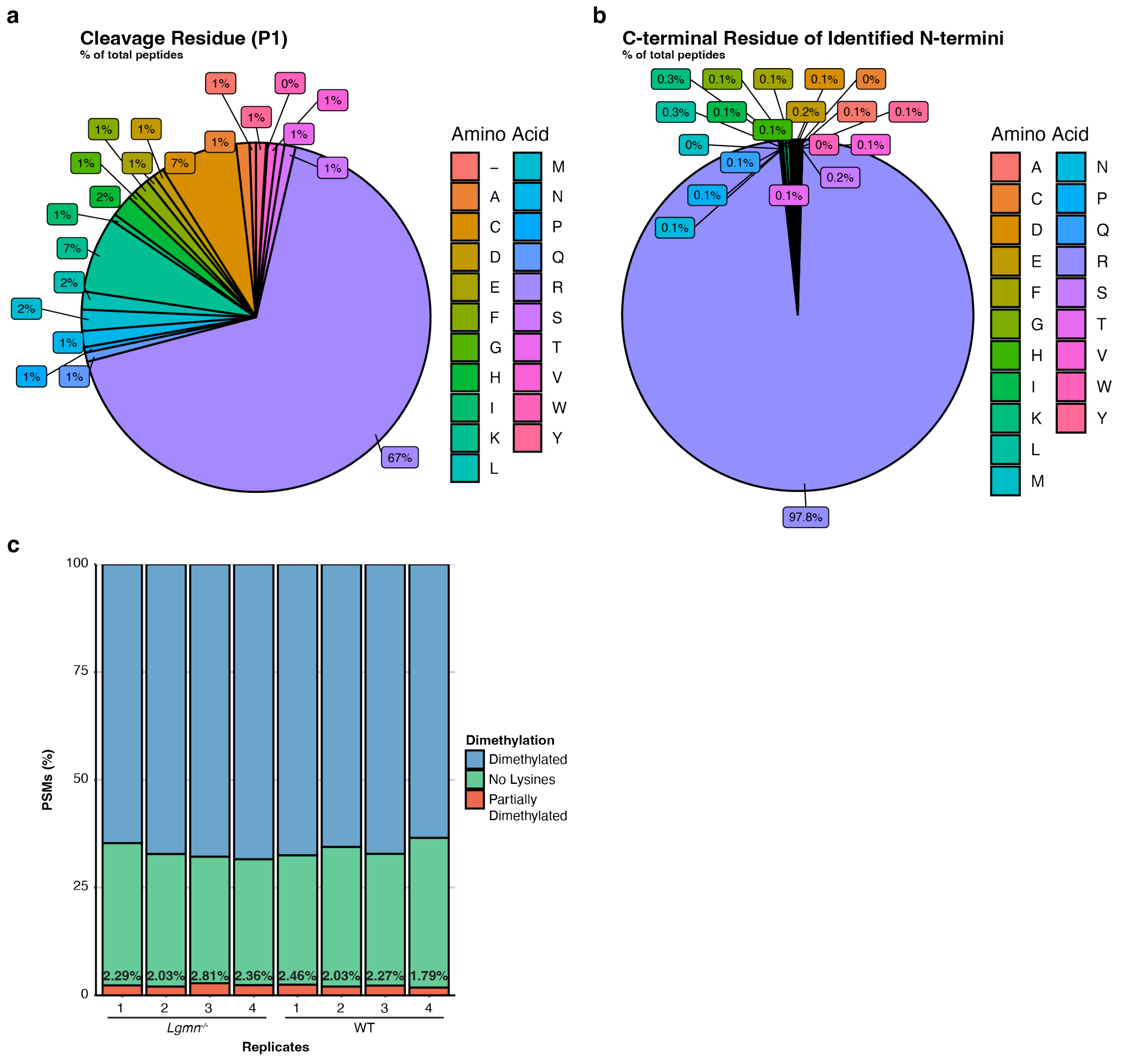


**Supplementary Fig. 9. Dimethylation efficacy of wildtype (WT) and legumain-deficient (*Lgmn*$^{-/-}$) naïve mouse spleens. a-b.** Spleen lysates were denatured, reduced, and alkylated prior to N-terminal dimethylation by formaldehyde. Following LC-MS/MS analysis, amino acid residues prior to the identified peptide/P1 residue (**a**) and at the end of each identified peptide (**b**) were used as measures of dimethylation efficacy. **c.** Dimethylation status of each peptide was also analysed according to whether all lysines were dimethylated (blue), no lysines were present (green), or lysines were partially dimethylated (red) for each biological replicate.

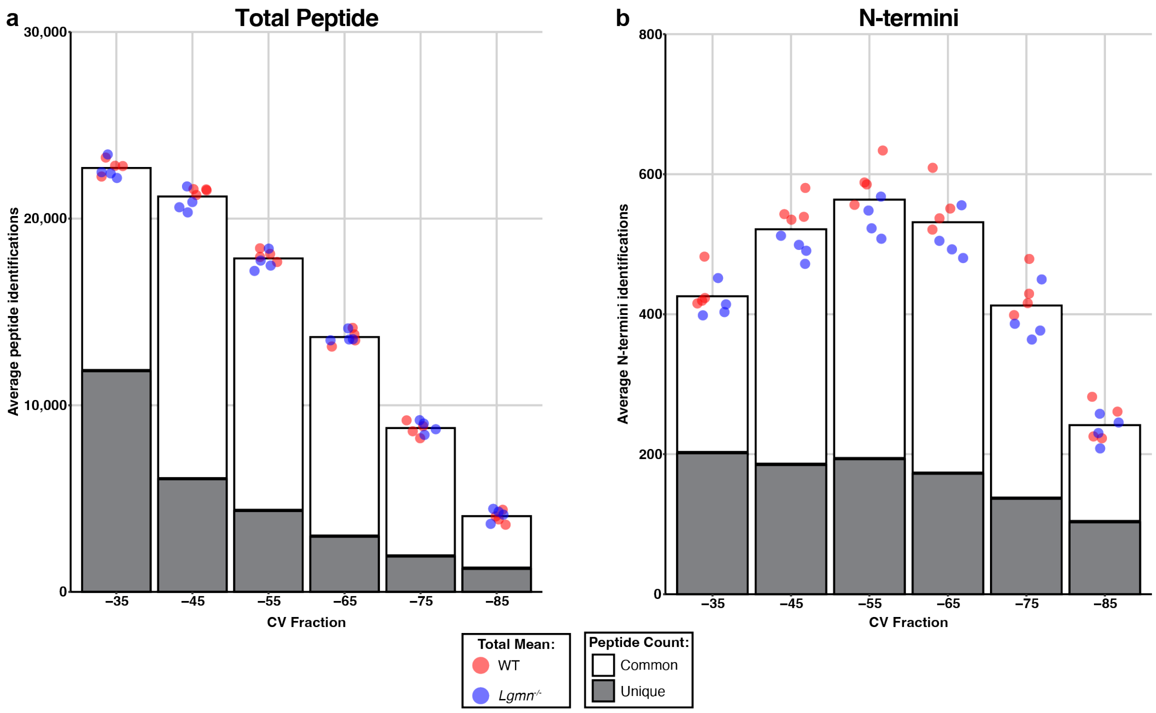


**Supplementary Fig. 10. FAIMS fractionation enables detection of unique peptides in each fraction. a-b.** Spleen lysates were prepared for mass spectrometry analysis on an Orbitrap 480™ mass spectrometer coupled to a FAIMS (high-field asymmetric waveform ion mobility spectrometry) device. Each biological replicate was fractionated into six individual fractions based on compensational voltage (CV) of -35, -45, -55, -65, -75, or -85. Following peptide-database matching using MSFragger (Fragpipe v.18.0), average number of identifications per biological replicate was visualised at the total peptide (**a**) and N-termini (**b**) level (n =4/group). Peptides and N-termini identified in only one CV fraction are shown in grey (unique), whilst those identified in more than one CV fraction are shown in white (common). Data points represent biological replicates where red indicates wildtype (WT and blue indicates legumain-deficient (*Lgmn*$^{-/-}$) spleen lysates.

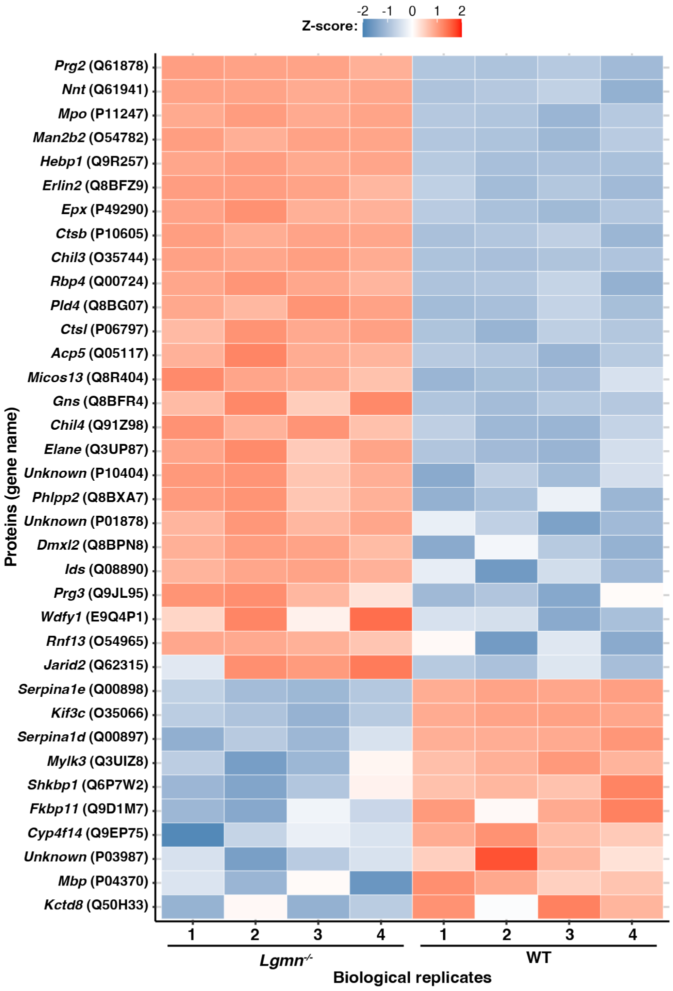


**Supplementary Fig. 11. Protein abundance changes observed in wild-type (WT) and legumain-deficient (*Lgmn*$^{-/-}$) spleen lysates are consistent across biological replicates.** Significantly enriched proteins (abs ($\log_2$(*Lgmn*$^{-/-}$/WT)) > 1 and -log10(p) > 2) were visualised on a heatmap. Z-scores were calculated based on the max label-free quantification (LFQ) intensity for each protein. Red indicates increased intensity from the mean and blue indicates decreased intensity from the mean. See Supplementary Table 15 for the complete list.

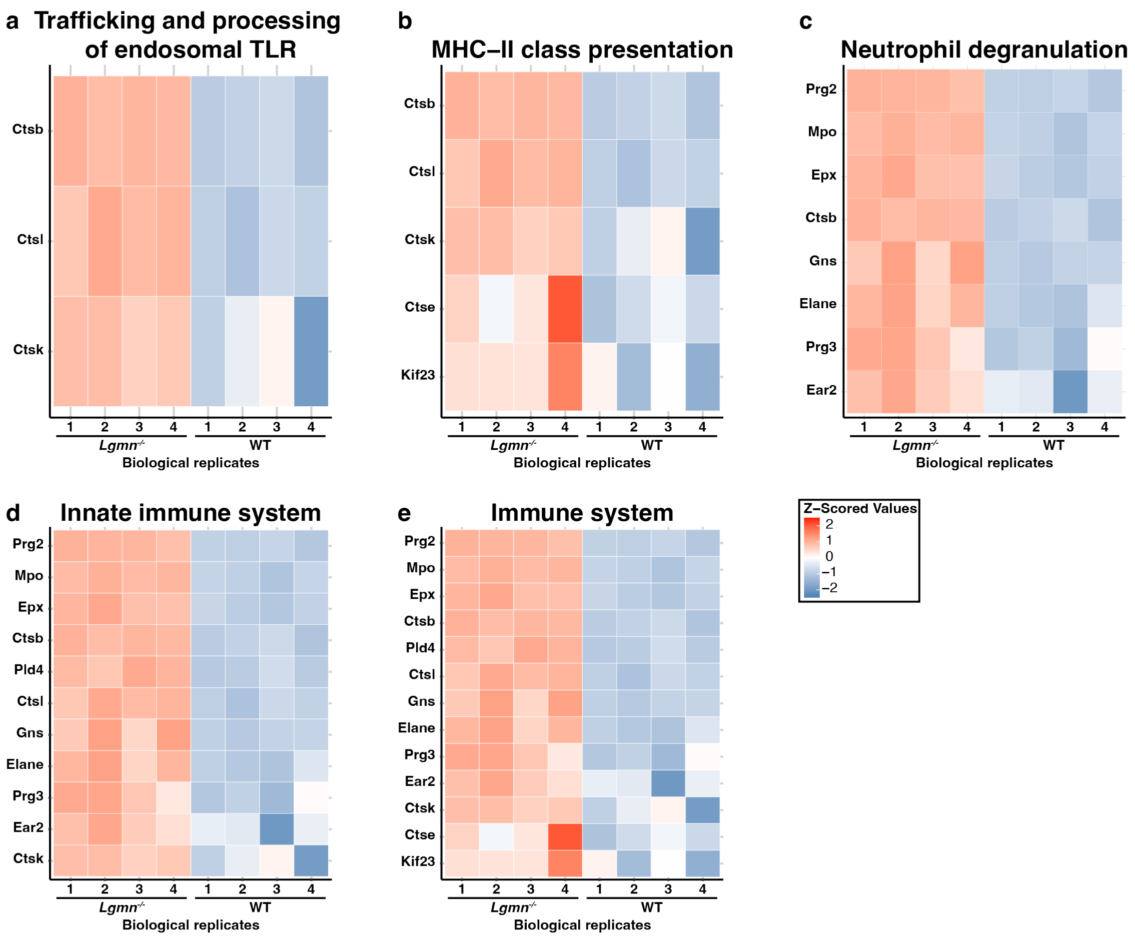


**Supplementary Fig. 12. Reactome pathway proteins are consistently upregulated in legumain-deficient (*Lgmn*$^{-/-}$) spleen lysates compared to wild-type (WT). a-e.** Following LC-MS/MS analysis of spleen lysates (n = 4/group), data were searched against an unreviewed mouse database in MSFragger (Fragpipe v.18.0) and quantified. A student's two-way t-test was performed in Perseus (v.1.6.0.7) to determine significant protein abundance changes between *Lgmn*$^{-/-}$ and WT spleen lysates. Proteins significantly upregulated in *Lgmn*$^{-/-}$ spleens were further analysed by STRING-dp for reactome pathways. Proteins included in each identified reactome pathway were extracted and max label-free quantification values for each biological replicate were used to calculate the z-score and deviation of each replicate from the mean (red = increased, blue = decreased). These values for each of the proteins are visualised as heatmaps.

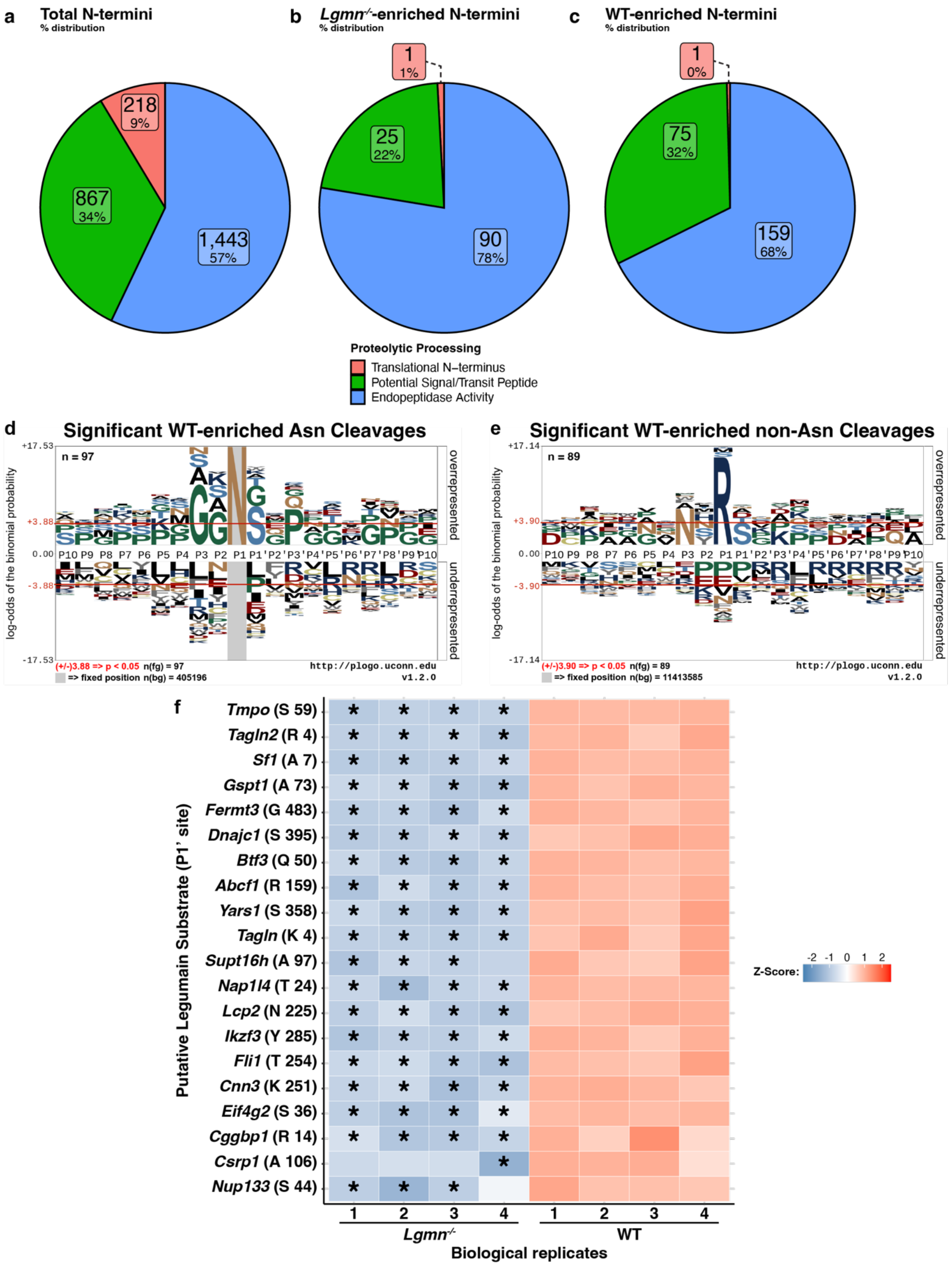


**Supplementary Fig. 13. Proteolytic processing in murine spleens is mainly a result of endopeptidase activity. a-c.** Following LC-MS/MS analysis of FAIMS-fractionated spleen lysates (n = 4/group), each identified N-termini was separated based on the position of the identified cleavage site in the protein where red indicates the translation N-terminus (1-2 aa), green indicates potential signal or transit peptides (3-65 aa) and blue indicates endopeptidase activity (66+ aa). **d-e.** Sequence motifs of N-termini significantly enriched in WT spleen lysates with asparaginyl (n = 97, putative legumain substrates) (**d**) or non-asparaginyl (n = 90) (**e**)

cleavages were created using plogo (O'Shea et al. 2013). Overrepresented amino acids appear above and underrepresented below the x-axis ($p < 0.05$). **f.** Top 20 legumain substrates identified are represented as a heat map. Z-scores were calculated based on the max label-free quantification (LFQ) intensities for each N-terminus. Red indicates increased intensity from the mean and blue indicates decreased intensity from the mean. Gene names and the identified N-termini are shown. Values which were imputed from a normal distribution (σ-width = 0.3 and σ-downshift = -1.8) are indicated by an asterisk (*).

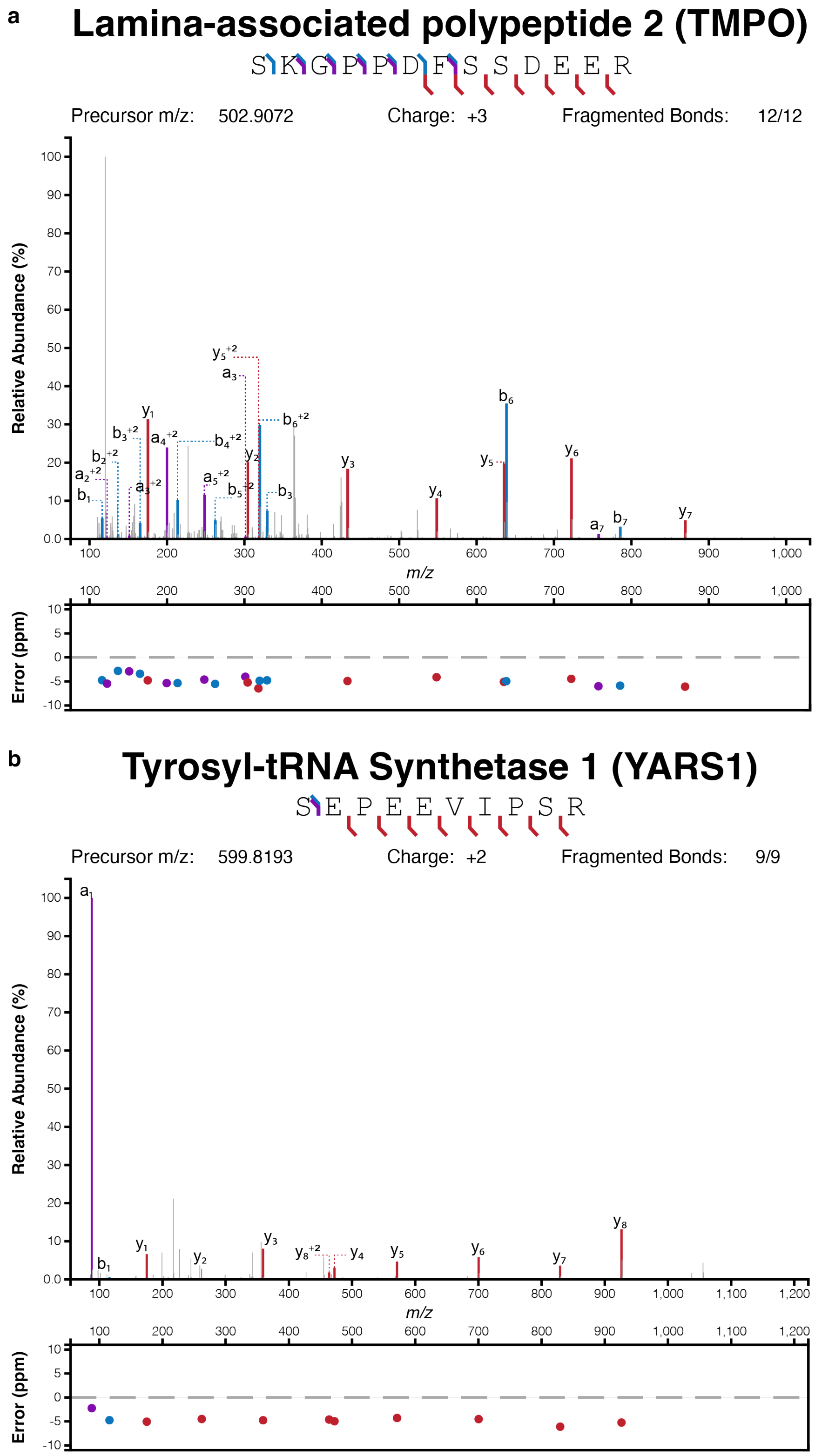


**Supplementary Fig. 14. Legumain directly processes various proteins *in vitro* as identified by FAIMS-enabled N-terminomics. a-b.** Recombinant proteins were incubated with activated recombinant legumain prior to N-terminomics analysis. MS2 analysis confirmed the dimethylation of the *Tmpo* peptide $^{59}$SKGPPDFSSDEER$^{71}$ and *Yars1* peptide $^{358}$SEPEEVIPSR$^{367}$ within legumain-treated samples supporting their cleavage. Figures were created using http://www.interactivepeptidespectralannotator.com/PeptideAnnotator.html with a fragment tolerance of ±10 ppm (Schmidt et al. 2021).
